# *In vivo* assessment of kinematic relationships for epithelial morphogenesis

**DOI:** 10.1101/2025.02.11.637562

**Authors:** Toshinori Namba, Kaoru Sugimura, Shuji Ishihara

**Affiliations:** Department of Integrated Sciences, Graduate School of Arts and Sciences, The University of Tokyo, Tokyo 153-8902, Japan; Universal Biology Institute, The University of Tokyo, Tokyo 113-0033, Japan; Department of Biological Sciences, Graduate School of Science, The University of Tokyo, Tokyo 113-0032, Japan; Department of Computational Biology and Medical Sciences, Graduate School of Frontier Sciences, The University of Tokyo, Chiba 277-8561, Japan

## Abstract

Tissue growth and deformation result from the combined effects of various cellular events, including cell shape change, cell rearrangement, cell division, and cell death. Resolving and integrating these cellular events is essential for understanding the coordination of tissue-scale growth and deformation by individual cellular behaviors that are critical for morphogenesis, wound healing, and other collective cellular phenomena. For epithelial tissues composed of tightly connected cells, the texture tensor method provides a unified framework for quantifying tissue and cell strains by tracking individual cells in live imaging data. The corresponding kinematic relationships have been introduced in a hydrodynamic model that we previously reported. In this study, we quantitatively evaluated the kinematic equations proposed in the hydrodynamic model using experimental data from a growing *Drosophila* wing. To accomplish this, we introduced modified definitions of the texture tensor and confirmed that one of these modifications more accurately represents approximated cellular shapes without relying on *ad hoc* scaling factors. By utilizing the modified tensor, we demonstrated the compatibility of the strain rate tensors and the accuracy of both the kinematic and cell number density equations. These results cross-validate the modified texture analysis and the hydrodynamic model. Furthermore, the precision of the kinematic relationships achieved in this study provides a robust foundation for more advanced integration of modeling and experiment.

## 1 Introduction

Morphogenesis is a self-organizing process that establishes multicellular bodies through the growth and deformation of tissues and organs. Tissue growth and deformation are driven by the behaviors of individual cells, which are coordinated through chemical and mechanical interactions among them [1–4]. Therefore, bridging the dynamics at tissue- and cell-scales is essential for understanding how precise and robust morphogenesis proceeds [4–6].

Epithelial tissues serve as excellent model systems for studying the mechanical control of morphogenesis because their relatively simple geometry and role in covering the body/organ surface make them accessible to live imaging and force/stress measurement [7–9]. Since epithelial cells are tightly connected and forces are transmitted along the plane of adherence junctions, a monolayer epithelium can be approximated as a two-dimensional tile of polygonal cells [10–12]. In such cohesive systems, tissue strain can be decomposed into strains resulting from various morphogenetic cell events, such as changes in cell shape, relative positions of cells (cell rearrangement), and cell numbers (cell division, and cell death or apoptosis) (Fig. 1a). Mathematical frameworks have been developed to quantify this decomposition of tissue strain into morphogenetic cell events (Fig. 1b, c) [13–17]. These strains were quantified as tensor variables and calculated from the experimentally observed cell geometry, providing a unified framework for quantifying strains resulting from various morphogenetic cell events, all within the same physical dimension. Along with strain measurement methods, corresponding continuum models have also been developed by coarse-graining cell-scale strains and stresses using a hydrodynamic description (Fig. 1c) [18–28]. These measurement and modeling techniques have begun to reveal the multi-scale integration in morphogenesis, specifically how cell-scale morphogenetic events and mechanical properties contribute to tissue-scale growth and deformation [14, 15, 29–32].

**Figure 1.**
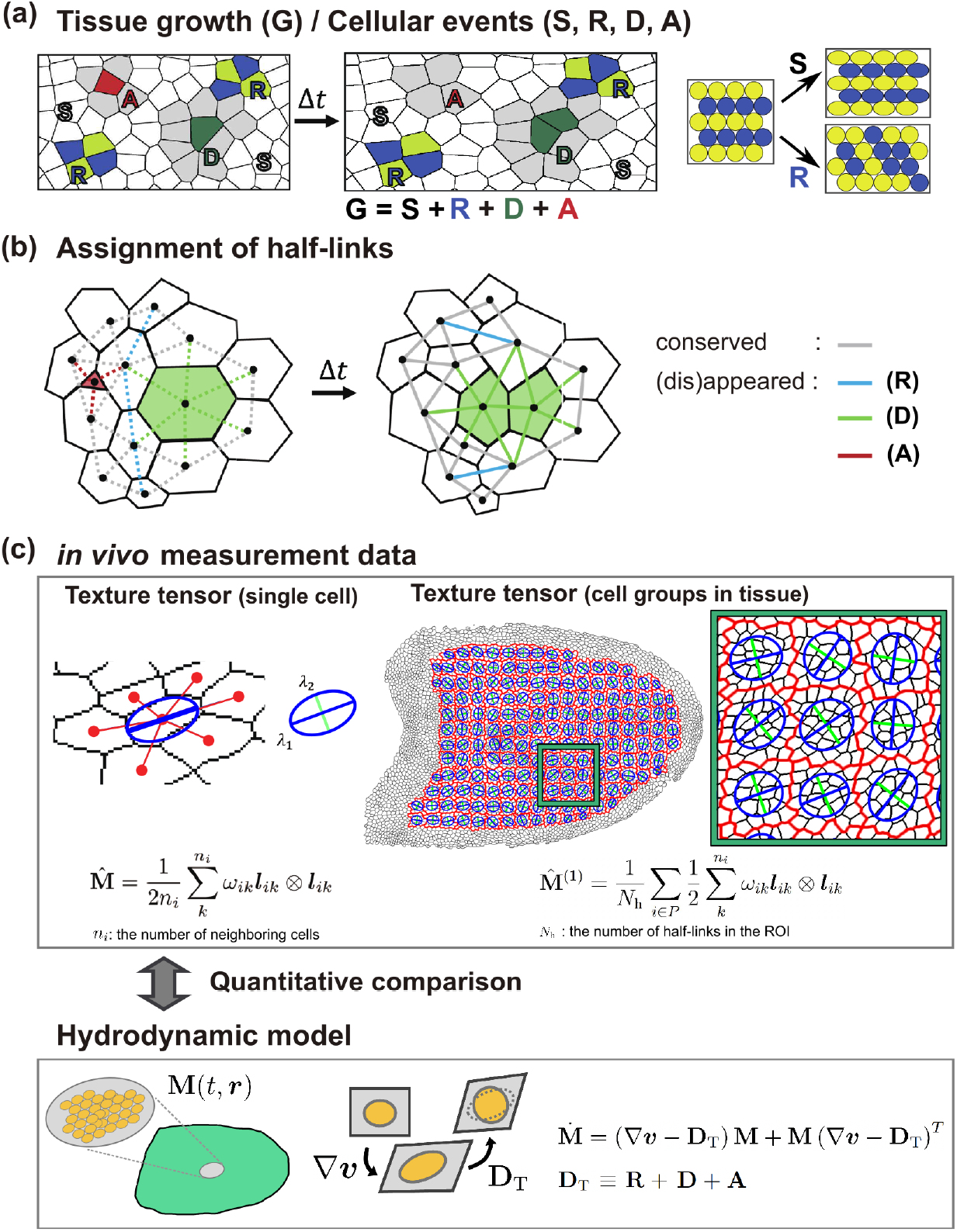
Quantification and kinematics of tissue deformation and morphogenetic cell events in epithelial tissue. (a) Morphogenetic cell events in a deforming epithelial tissue. The strain tensor **G** represents horizontal elongation of tissue, where its constitutive cells undergo shape change (white; S), rearrangement (blue and yellow; R), division (green; D), and apoptosis (red; A). Division and apoptosis induce the deformation of surrounding cells (gray). Tissue deformation is the sum of strains generated by these cellular events. On the right, examples of tissue deformation driven solely by cell shape change or cell rearrangement are illustrated. (b) Assignment of cell-to-cell half-links to each morphogenetic event. The links corresponding to each event are color-coded as shown on the right. (c) Quantification (top) and kinematic equations (bottom) of tissue deformation and morphogenetic cell events. Top: The major (blue) and minor (green) axes of ellipses correspond to the eigenvectors of the texture tensor ― computed for either a single cell (left; 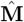) or a group of cells (right;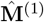). Red outlines denote the boundaries of regions of interest (ROIs) used to compute the coarse-grained texture tensor 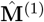. See also Fig. S1 for the data analysis workflow. Bottom: Kinematic equations in the multi-scale continuum model for epithelial mechanics [19] indicate that the time evolution of the cell shape tensor **M** results from the tissue strain rate ∇***v*** and the topological strain **D**_T_ = **R** + **D** + **A**.

A quantitative comparison of the aforementioned hydrodynamic models with *in vivo* data on cell mechanics will serve to validate the models and establish a basis for a more advanced integration of modeling and experiments. The hydrodynamic description of material deformation generally includes kinematics, kinetics, and force balance [33–35]. Kinematics describes the geometric relationships governing the time evolution of deforming objects, whereas kinetics represents the mechanical properties of materials, such as constitutive equations. Kinetics are often more intricate, involving rheology, *that is*, elasticity, viscosity/plasticity, and potentially active forces, and are often tissue-specific. In contrast, kinematic relationships provide more general and robust frameworks characterized by a few parameters, making them applicable across different epithelial tissues. Therefore, when assessing continuum models against experimental data, the kinematic relationships proposed in the theory should first be assessed.

The primary objective of this study is to quantitatively assess the kinematic equations proposed in the hydrodynamic model for epithelial mechanics [18,19] by analyzing experimental data obtained from a developing *Drosophila* wing. To achieve this, we utilized the unified quantification method known as texture tensor analysis to measure tissue and cell strains [14]. However, the original definition of the texture tensor, used to represent the cell shape, may not be optimal for a direct comparison between the model and experimental data owing to its lack of normalization with respect to cell geometrical quantities. To address this limitation, we introduced a modified definition of the texture tensor and evaluated its effectiveness in approximating cell shape by assessing its ability to capture cell area and second moments. We also examined the compatibility of the strain rate tensor derived from texture tensor analysis with that calculated from the symmetric part of the strain tensor, as measured using particle image velocimetry (PIV), a technique used to evaluate the velocity field from images. The agreement between these two methods is a prerequisite for validating tensor analysis, but it had not been explicitly demonstrated in previous studies. By applying the modified tensor analysis to *Drosophila* pupal wing data, we demonstrated the accuracy of the kinematic equations proposed in [18] and [19]. This, in turn, supports the validity of the tensor analysis using the modified texture tensor.

The remainder of this paper is organized as follows: Sec. 2 outlines the kinematics of tissue deformation and clarifies the purpose and problems addressed in this study. Theoretical and experimental formulations involving various tensors, such as the strain rate and cell-shape tensors, are introduced. Sec. 3 describes the materials and methods utilized in this study. Sec. 4 presents the analysis results, including the introduction and evaluation of a new definition of the texture tensor, assessment of cell number density, and kinematic equations. Finally, Sec. 5 summarizes the findings and discusses their implications.

## 2 Multi-scale hydrodynamic model and data analyses for epithelial mechanics

This section introduces kinematics in a multi-scale hydrodynamic model (Section 2.1 and 2.2) [19] and a related data analysis framework based on the texture tensor (Section 2.3) [14, 36]. Kinematic equations to be tested on experimental data are presented, along with an explanation of the methods utilized to measure kinematics from time-lapse movies of epithelial tissue.

### 2.1 Strain rate tensors for tissue deformation and accompanying morphogenetic cell processes

Tissue deformation as a two-dimensional material is described by the strain rate tensor

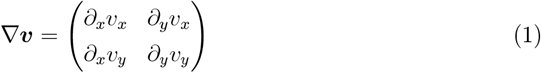

where ***v***(*t*, ***r***) represents the velocity field associated with tissue deformation at time *t* and position ***r*** = (*x, y*) (note that some literature used a transposed definition for ∇***v*** [13,36,37]). The strain rate tensor is decomposed into the symmetric and asymmetric parts. The symmetric part is given by

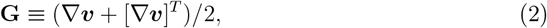

where the superscript ^*T*^ denotes the transpose of the tensor and the asymmetric part **Ω** ≡ (∇***v*** − [∇***v***]^*T*^ )*/*2 represents rotation. Because material deformation can only be observed at discrete time intervals, data analysis should relate the strain tensor ∇***v*** to the deformation that occurs during the time interval Δ*t*. Let the point ***r*** at time *t* move to ***R*** at time *t* + Δ*t*. The deformation gradient tensor **F** between the consecutive time points is defined as follows:

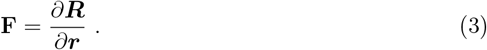

Note that ***R***, and therefore **F**, is a functions of *t* and ***r***. For a sufficiently small Δ*t*, **F** is related to the strain tensor Eq. 1 as ∇***v*** = (**F** − **I**)*/*Δ*t* with a 2 *×* 2 identity tensor **I**. The symmetric part of the strain rate tensor **G** (Eq. 2) can be approximated as:

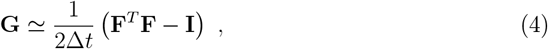

which follows from substituting the relation **F** = **I**+∇***v***Δ*t*. This approximation becomes exact in the limit Δ*t* → 0.

Tissue deformation results from morphogenetic cellular events, including cell shape change (S), rearrangements (R), divisions (D), and apoptosis (A) (Fig. 1a). The tissue strain rate **G** can be decomposed into the strains associated with each of these cellular events as follows:

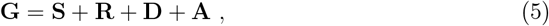

where **S, R, D**, and **A** represent the symmetric strain rate tensors resulting from cell shape change, rearrangement, division, and apoptosis, respectively. In the multi-scale continuum theory, the strain **S** is associated with the resilient force responsible for maintaining cellular shape, while **R, D**, and **A** represent strains that accompany changes in relative cellular positions and do not contribute to resilient forces. Thus, we refer to **G** and **S** as total and elastic strains, respectively, while classifying the others as plastic strains.

### 2.2 Kinematic equations for two-dimensional cell tiles

The kinematic equations for a two-dimensional tile of epithelial cells was proposed as a coarse-grained representation of the deformation resulting from morphogenetic cell events [18]. Under the assumption that the cellular shape can be approximated by an ellipse, it is quantified using a 2 *×* 2 matrix **M** [36]. The matrix **M** is symmetric and positive-definite, and has dimensions of squared length. The two eigenvectors of **M** point to the longer and shorter axes of the ellipse, with the corresponding eigenvalues representing the squared lengths of the radii. The cell area is approximated from the tensor **M** as follows:

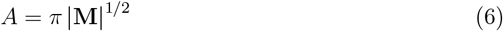

where |**M**| denotes the determinant of **M**. For a continuum description of multicellular tissues, the tensor field variable **M**(*t*, ***r***) is introduced as a measure of the coarse-grained cell shape at time *t* and position ***r*** (bottom panel in Fig. 1c). Because the epithelial tissue comprises a tile of cells, the evolution of cell shape over time is closely linked to tissue deformation, quantified by the velocity field ***v***(*t*, ***r***). Theoretical analysis led to the relationship [18, 19]

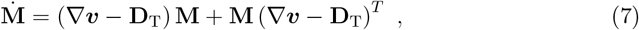

where 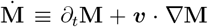 represents the Lagrange derivative of **M** and **D**_T_ ≡ **R** + **D** + **A** = **G** − **S** denotes a strain rate tensor resulting from morphogenetic cell events accompanied by topological changes in cell-neighbor relationships. This equation is hereafter referred to as the kinematic equation for **M**. The equation comprises no mechanical parameters and must be satisfied if the tensors **M** and **D**_T_ are appropriately defined and measured from the image data.

Cell number density changes owing to alterations in cell shape, division rates, influx, and efflux. Cell number density is simply the inverse of cell area, expressed by *ρ* = 1*/A* = 1*/π*|**M**|^1*/*2^. Using the identity *d*|**a**|*/dt* = |**a**|Tr (**a**^−1^*d***a***/dt*) for an arbitrary regular matrix **a** that is a function of a variable *t*, the equation for changes in cell number density is derived from Eq. 7 as follows:

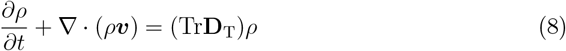

where Tr**D**_T_ denotes the trace of **D**_T_. This equation represents the cell number conservation over time, where Tr**D**_T_ corresponds to the variation rate of cell number density. Similar to Eq. 7, the equation contains no mechanical parameters and should be satisfied accordingly.

The trace of the strain rate tensor ∇***v*** represents the local rate of area change (*i*.*e*., expansion or contraction) at that point. Because cell rearrangement does not contribute to area changes, the following assumption should apply:

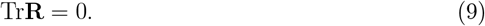

This indicates that **R** is not involved in the variation rate of cell number density. Traces of cell division and apoptosis tensors, **D** and **A**, correspond to cell division and death rates, respectively, with anticipated values Tr**D** ≥ 0 and Tr**A** ≤ 0.

Notably, the derivation of the cell number density equation in Eq. 8 is a natural outcome of the kinematic equation for **M** (Eq. 7) [18, 19]. This integration aligns with the characteristics of tissue, where changes in cell shape are inherently linked to tissue deformation (see also the discussion in Sec. 5).

### 2.3 Coarse-grained and unified quantification of morphogenetic cell events using the texture tensor

Experimental measurement of the quantities explained above is not a trivial problem, sfor which one needs to properly define the cellular shape and its temporal variation, and their decomposition in accordance with Eq. 5. The texture tensor, a 2*×*2 matrix representing coarse-grained cell shape, can be obtained from experimental data including links between the centers of neighboring cells (Fig. 1b, c). The texture tensor was originally defined in [14, 36] as follows:

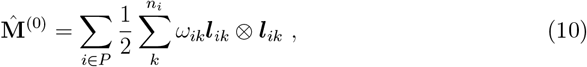

where, ***l***_*ik*_ = ***x***_*k*_ − ***x***_*i*_ denotes the vector connecting the cell centers of cell *i* and its neighboring cell *k*. The hat symbol 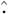 indicates a tensor derived from half-links ***l***_*ik*_ (this convention applies throughout this manuscript). The sum of *ω*_*ik*_***l***_*ik*_ ⊗ ***l***_*ik*_ is computed over cells in the region of interest (ROI) indexed by *P* . The sum over *k* indicates the sum over the links connecting cell *i* with its neighboring cells, where *n*_*i*_ represents the number of neighboring cells. The weighting parameter *ω*_*ik*_ is set to *ω*_*ik*_ = 1 if neighboring cells are in contact with the edge, and *ω*_*ik*_ = 1*/*2 if they are in contact with the four-fold vertex. Because each link was counted twice, a factor 1*/*2 is applied, each link ***l***_*ik*_ is thus designated as a half-link. The half-links to and from the outmost cells are excluded when calculating 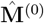.

By using the temporal change in the texture tensor 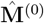, Guirao et al. [14] developed a mathematical framework for calculating the total strain **G** and the decomposition to **S, R, D**, and **A** to align with Eq. 5 (Fig. 1c and Fig. S1; Supporting Information A). In this framework, the deformation gradient tensor of ROI, denoted as 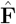, is computed from the difference in 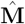 between consecutive time points *t* and *t* +Δ*t* (Fig. S1d). While this framework can accommodate data in both Eulerian and Lagrangian descriptions, our previous and current studies have employed Lagrangian tracking (*i*.*e*., the individual cells are tracked from the initial to the final time points) [14]. This approach allows us to define ROIs with one-to-one correspondence across time frames (Sec. 3.7), enabling the evaluation of temporal changes in physical quantities between time points *t* and *t* + Δ*t*, while excluding the effects of influx and efflux, and facilitating the computation of their Lagrange derivatives (see Sec. 3.8).

Subsequently, the symmetric part of the strain tensor, 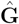, is determined using Eq. 4, as follows:

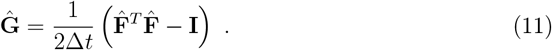

Simultaneously, by tracking the appearance and disappearance of half-links, the decomposition of 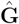 satisfy into 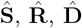, and 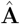 are computed from cellular contour images to satisfy

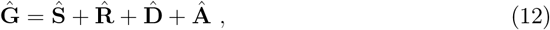

corresponding to Eq. 5.

The details of this method are summarized in Supporting Information A and B. Our method has been slightly modified from the original to more directly calculate 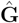 , with the difference remaining small―on the order of *𝒪* (Δ*t*^2^) (Supporting Information A). Additionally, extracting strain rate tensors from experimental data involves different assignment options, even when skeletonized images are provided. The original assignment rules [14] were validated as outlined in detail (Supporting Information B).

While the method requires cell contour data from time-lapse images to compute 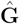 and its decomposition 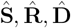 and 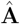, the strain rate tensor **G** can also be determined from the velocity field of a tissue, which can be obtained using less intensive image processing methods such as PIV. Comparison of strain rate tensors obtained by the two different measurement is performed in Sec. 4.3.

## 3 Materials and Methods

### 3.1 *Drosophila* genetics

*Drosophila* stocks were maintained at 25^*°*^C. The genotype of the three wild-type samples utilized in this study is DE-cad-GFP [38].

### 3.2 Image acquisition

The preparation of *Drosophila* pupal wing samples for image collection was conducted following previously established protocols [14, 39]. Briefly, pupae at appropriate developmental stages were dissected to remove the pupal case covering the left wing and were then positioned on a small drop of Immersol W 2010 (Zeiss 444969-0000-000) in a glass bottom dish. Images were acquired at 25^*°*^C using an inverted confocal spinning disk microscope (Olympus IX83 combined with Yokogawa CSU-W1) equipped with an iXon3 888 EMCCD camera (Andor), an Olympus 60x/NA1.2 SPlanApo water-immersion objective, and a temperature control chamber (TOKAI HIT), utilizing IQ 2.9.1 (Andor) [14].

### 3.3 Image data

Two types of image data were analyzed (Fig. 2a): 1. Whole-wing images acquired at 5-minute intervals starting from 15.5 hours after puparium formation (h APF). 2. C-region images acquired at 1-minute intervals starting from 24 h APF. One of the three former type images was published in [14], and all three of the latter type images were examined in [40].

**Figure 2.**
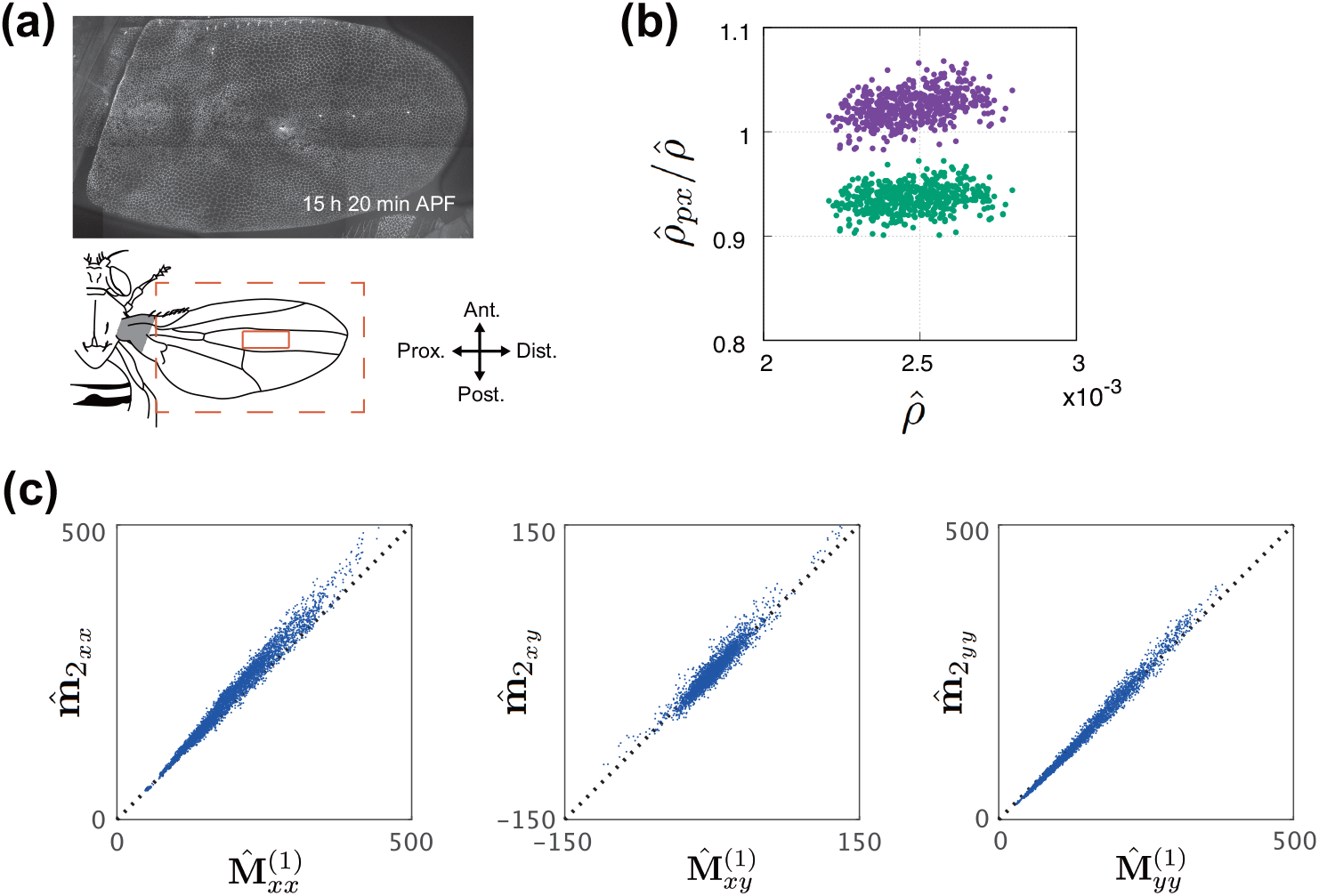
Validation of the texture tensor 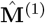 as a cell shape indicator. (a) Top: Image of DE-cad-GFP from the *Drosophila* pupal wing at 15 h 20 min APF. Bottom: Schematic of the *Drosophila* pupal wing. We analyzed images of the whole-wing (black orange square) and the C region (orange square). (b) Ratio of cell number density measured by direct pixel counting 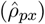 to that measured by the texture tensor 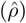 is shown. Direct pixel counting of 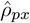 was performed both excluding the outline pixels (purple points) and including half of the outline pixels (green points). Each data point represents the ratio value for a single ROI in the C region. (c) Comparison between the texture tensor 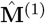 (Eq. 13) and the mean second moment of cells, 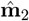 (Eq. 15). Each component of these tensors is indicated in the respective panels.

### 3.4 Image processing

Image segmentation and tiling were performed using custom-made macros and plug-ins in ImageJ [14,41]. Vertex position and connectivity were extracted from the skeletonized images using custom-made code in OpenCV [41]. See also Fig. S1 for the data analysis workflow.

### 3.5 Particle Image Velocimetry (PIV)

The displacement within sequential images was measured using **PIVlab**, an open-source tool in MATLAB [42], with a correlation algorithm *FFT window deformation* (direct Fourier transform correlation with multiple passes and deforming windows), along with ensemble correlation. The interrogation window and window steps were set at 64*×*64 and 16 pixels, respectively. Erroneous vectors were eliminated and supplemented by the standard deviation filter with a parameter of *n* = 3.

### 3.6 Cell tracking

The cell IDs were assigned by labeling each segmented area of the skeletonized images using the function **bwlabeln** in MATLAB. From the assigned cell IDs, the IDs of all neighboring cells were obtained, with cells adjacent to the outer region designated as the outermost cells. Subsequently, a cell tracking procedure was implemented. Initially, the *x*-*y* coordinates of the cell center were adjusted based on the vector field measured using PIV. Second, a cost matrix, defined by the square Euclidean distance between cell centers, was optimized using the function **assignjv** in MATLAB to determine the correspondence of cell IDs in two consecutive time frames [43]. To reduce the computational cost, the components of the cost matrix for cell pairs over 400 pixels apart were set to infinity. Third, cell division and apoptosis were detected based on the appearance and disappearance of cells, respectively. We examined the changes in the area of neighboring cells surrounding newly emerged cells, designating the cell with the largest decrease in area as a dividing cell. While this method may not be effective in cases where neighboring cells undergo division and apoptosis simultaneously, such occurrences were rare in the time-lapse images analyzed in this study. We thoroughly reviewed all cell tracking patterns and made manual adjustments as needed.

### 3.7 Setting ROI

To calculate texture tensors and strain rates from the time-lapse movies of the pupal wing, the image at the initial time point, either 15.5 h APF (whole-wing movie) or at 24 h APF (C region movie), was divided into 120*×*120 pixel tiles. ROIs were then defined as regions composed of cells approximating these rectangular tiles (Fig. S1d). Each ROI defined at the initial frame was tracked over time to determine the corresponding ROIs in subsequent frames. Thus, ROIs across different time points maintain mutual correspondence—that is, they are composed of the same cells, or of their mother or daughter cells. An ROI of this size contains approximately 30 cells at 15.5 h APF and approximately 60 cells at 32 h APF. Previous research has shown that this ROI size produces consistent patterns of tissue deformation, cellular events, and stress in the *Drosophila* pupal wing and notum [14, 44].

### 3.8 Computation of time derivative of coarse-grained quantities

Coarse-grained quantities are defined using the ROIs described in Sec.3.7. Temporal changes in physical quantities are calculated from corresponding ROIs between consecutive time points *t* and *t* + Δ*t* (Fig. S1c), allowing for the evaluation of Lagrange derivatives; for example, 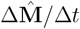 serves as a measure of the Lagrange derivative 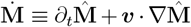 . For the calculation of time derivatives and the estimation of 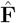, we assumed that the deformation within each ROI is sufficiently small and that most half-links are conserved between time points *t* and *t* + Δ*t*. In our analysis, a 5-minute time interval satisfies this assumption. As confirmed in the Results (Sec. 4), second-order terms in Δ*t* are negligible, supporting the validity of this approximation

### 3.9 Assignment of half-links to cellular events

We implemented the methodology outlined in Ref. [14] to assign each half-link to one of the cellular events, such as cell shape change, rearrangement, division, and apoptosis (Fig. 1b; refer to Supporting Information B for details).

## 4 Results

### 4.1 Modified definitions of the texture tensor

The original texture tensor, defined in Eq. 10, 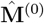 , exhibits a scaling behavior with the number of cells within the ROI, *N*_c_. This dependence on the choice of ROI is incompatible with the definition of cell shape tensor used in the hydrodynamic model (Sec. 2.2) [19], which is based on the coarse-grained representation of cell shapes and remains invariant to the number of cells within the ROI. We therefore introduced an alternative form of the texture tensor as follows:

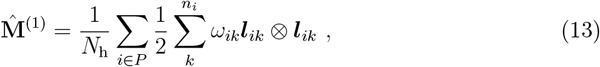

where *N*_h_ denotes the number of half-links in the ROI. In this definition the texture tensor is normalized by *N*_h_ rather than being proportional to *N*_c_. Furthermore, we also considered other possible forms of the texture tensor (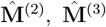 , and 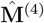, in Supporting Information C). All the proposed definitions of texture tensors possess a physical dimension of squared length, differing primarily in the normalization procedure based on the number of cells and their adjacent counterparts.

### 4.2 Evaluation of the cellular area and second moment using the modified texture tensors

To assess the accuracy of the modified texture tensors in capturing the cell morphology within epithelial tissues, we conducted a comparative analysis of the cellular area and second moment measurements obtained using different texture tensor formulations against those derived directly from pixel counting in image data of the *Drosophila* pupal wing (Fig. 2). The ratio of cell number density measured by direct pixel counting to that measured by the texture tensor 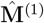 is shown in Fig. 2b, where the individual dots represent data from single ROIs of 120 pixels *×* 120 pixels, with averages computed across ROIs of this specific size, unless otherwise specified hereinafter. The ratio exceeded unity when excluding pixels along cell contours (Fig. 2b, magenta dots) but decreased when half of the pixels on the cell contours were included (Fig. 2b, green dots). These results suggest that the modified texture tensor 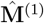 accurately measures the cell area, with deviations from unity attributed to image resolution in the measurement.

Next, we compared 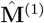 with the second moment of the cellular shape, quantified using pixel counting. The second moment of a cell, based on pixel data defining cellular shape, is calculated as follows:

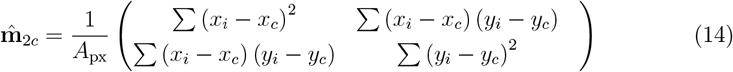

where the sum is taken over all pixels within the cell, with each pixel located at (*x*_*i*_, *y*_*i*_). The coordinates (*x*_*c*_, *y*_*c*_) represent the center of the cell (*e*.*g*.,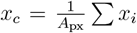 ) and *A*_px_ denotes the cell area as determined by pixel counting. We averaged the second moment of the cells over the ROI and calculated

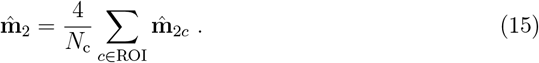

We applied a factor of 4 because an ellipse with radii *r*_*a*_ and *r*_*b*_ along the *x*- and *y*-axes has second moments 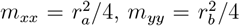, and *m*_*xy*_ = 0. Therefore, 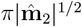 corresponds to the area, which is consistent with Eq. 6. As shown in Fig. 2c, the texture tensor 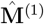 closely aligns with the pixel-based mean second moment of the cellular shape in each ROI, 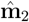. These findings collectively demonstrate that 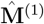 accurately measures cellular area and second moment from real image data.

For comparison, we assessed alternative definitions of the texture tensors against the second moments of the cellular shape, 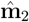. We plotted 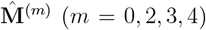 and 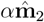 in Figs. S2a-d, where the scaling factor *α* is introduced as a fitting parameter determined by the mean value of the cell area. 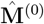 and 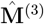 did not align with the second moment 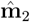 (Figs. S2a and c). This discrepancy stemmed from the fact that they were not normalized by the cell number *N*_c_, which varies across the dataset. This is in sharp contrast to 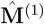, which only requires the predetermined factor of 4.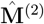 and 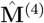 aligned significantly, comparable with 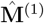. The scaling factor was *α* ∼ 5.8 for 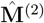, which can be attributed to the fact that 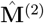 differed from 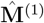 only in the scaling ratio *N*_h_*/N*_c_, which is expected to be close to 6 (Fig. S2b). 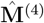 showed results almost identical to 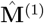 with a similar scaling factor *α* = 0.95 (Fig. S2d).

### 4.3 Consistency in the strain rate measured using 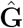 and PIV

In Sec. 4.2, we validated the utilization of the modified texture tensor 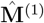 to measure the morphology of cells using still images. Below, we assessed its ability to measure the dynamic deformation of tissues. The deformation gradient tensor 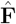 and the symmetric part of the strain rate tensor 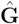 were calculated from the time variation of the texture tensor, which requires segmented images that highlight cell contours. However, the strain rate tensor ∇***v*** can also be determined by PIV without image segmentation, providing a direct characterization of tissue deformation. These two quantities 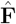 and ∇***v*** should be related to 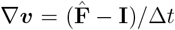, assuming a sufficiently small time interval Δ*t*. Ensuring consistency between the left- and right-hand sides obtained using different measurement methods is a prerequisite for validating the analysis based on the texture tensor.

As shown in Fig. 3a, (∇***v***)Δ*t* measured using PIV and 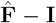 calculated from the texture tensor 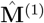 agreed with each other (see Fig. S3 for results from different samples of whole-wing image data). Moreover, when Δ*t* is small, 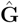 can be approximated by the symmetric part of ∇***v*** (Eq. 2), a relationship validated in Fig. 3b and Figs. S4. In our dataset, Δ*t* = 5 min, the relative magnitude of the discrepancy between Eq. 11 and Eq. 2 is ∼ 10^−2^ times smaller than that of the symmetric term, which makes it negligible. These findings validate 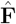 and 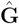 derived from 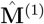 as indicators of dynamic tissue deformation.

**Figure 3.**
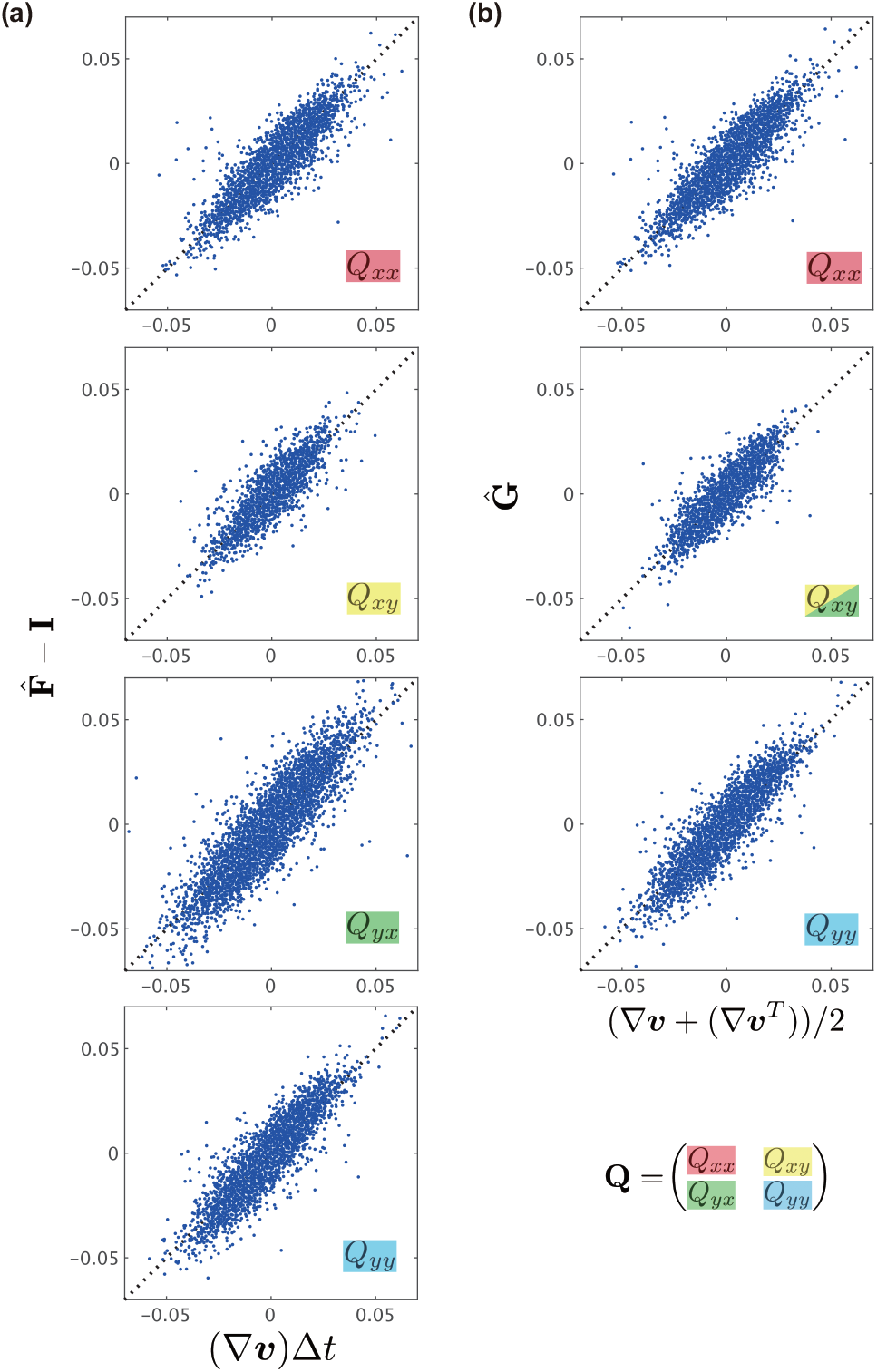
Validation of the strain rate measurement. (a) Comparison of the strain rate tensor ∇***v*** and deformation gradient tensor 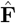. The average of the vectors obtained by PIV over the ROI was used to compute the strain rate tensor ∇***v***. (b) Comparison of the components of 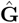 with the symmetric part of ∇***v*** obtained from PIV.

### 4.4 Assessment of the cell number density equation

Having confirmed the applicability of the modified texture tensor 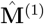 for both static cell shape analysis (Sec. 4.2) and dynamic deformation analysis (Sec. 4.3), we next investigated whether the time evolution equation for cell number density (Eq. 8) in our continuum model aligned with the *in vivo* data obtained using 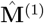.

In Eq. 8, Tr**D**_T_ = Tr**R**+Tr**D**+Tr**A** represents the change in the cell number density, with **R, D**, and **A** denoting the strains associated with cell rearrangement, division, and apoptosis, respectively. Given that cell rearrangement does not alter tissue area, Tr**R** is expected to be zero (Eq. 9), whereas cell division and apoptosis contribute to the area expansion (Tr**D** ≥ 0) and contraction (Tr**A** ≤ 0), respectively. We examined whether the data from the wings exhibited the anticipated features (Fig. 4a). Our analysis of wing data revealed that 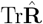 was significantly smaller than the other components throughout the time course of wing development, supporting the relationship Tr**R** = 0 (Eq. 9). Furthermore, Tr**D** was consistently positive, as expected, whereas 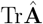 was nearly zero (∼ 1.5*×*10^−6^h^−1^), indicating that apoptosis contributes minimally to deformation of the pupal wing. These results are consistent with the low frequency of apoptosis reported in the wing [14]. The findings serve as a foundation for validating the cell number density equation outlined below.

**Figure 4.**
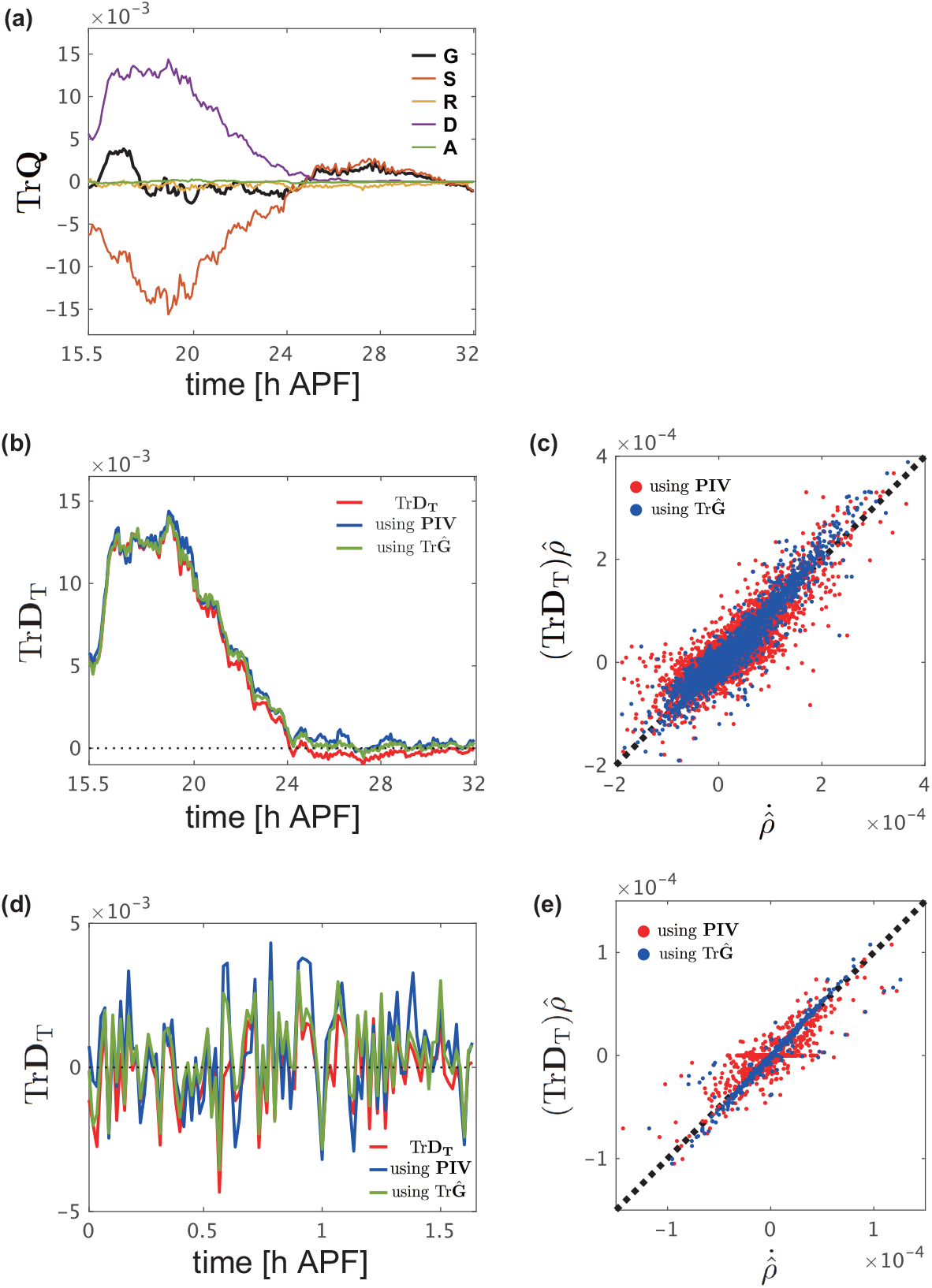
Assessment of the cell number density equation. (a) Time series of the traces of 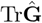 (black), Tr**Ŝ** (red), 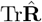 (yellow), 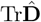 (purple), and 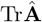 (green) for time-lapse data of the whole-wing captured at 5-min intervals starting from 15 h 30 min APF. (b, d) Time series data indicating the value obtained by dividing the value on the left-hand side of Eq. 8 by 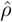 (blue and green lines), compared with Tr**D**_T_ (red line), for the time-lapse data of the whole-wing captured at 5-min intervals starting from 15 h 30 min APF (b) and time-lapse data of the C region of the wing captured at 1-min intervals starting from 24 h APF (d). Blue line: The value obtained by using the velocity field measured using PIV for ∇ ·***v***. Green line: The value obtained by using the trace of the strain rate tensor (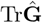) for ∇ · ***v***. These time series data denote the moving average values with a window of width 30 min (b) and 3 min (d). (c, e) Comparison of Eq. 8 using different measurement methods of ∇ · ***v***. The analysis results are shown using the velocity field measured by PIV (red dots) and the trace of the strain rate tensor (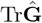, blue dots). Each point represents data from an individual ROI. Data analyzed in (b) and (d) are used in (c) and (e), respectively.

Next, we evaluated the cell number density equation (Eq. 8) by employing different measurement methods for tissue deformation and assessing the consistency among them. Specifically, the left-hand side of Eq. 8 was evaluated as follows: We separated the left-hand side as 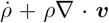, where 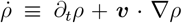. The time derivatives for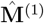, and consequently for the density 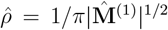, were calculated as the difference between two consecutive time frames. In this way, we evaluated the Lagrange derivative 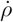 (but not *∂*_*t*_*ρ*), since the variables were measured using ROIs defined by tracking individual cells, as described in the Materials and Methods section 3.8. Furthermore, the divergence of the velocity field, *∇ ·* ***v***, can be quantified in two ways; one using the total tissue strain rate **G** obtained using the texture tensor through *∇ ·* ***v*** = Tr**G**, and the other by direct velocity field measurement using PIV. The right-hand side Tr**D**_T_ was quantified by summing the respective contributions from morphogenetic cell events, specifically 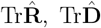, and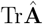. Collectively, there are three ways to calculate the temporal evolution of the cell number density: 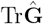 and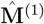, PIV and 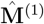, and the sum of the decomposed Tr**D**_T_. The time series data obtained from these measurements were plotted, with averages presented for the whole-wing (Fig. 4b) and the C region of the wing (Fig. 4d). The results from the two measurements of the velocity divergence *∇ ·* ***v*** were almost identical (blue and red lines in Figs. 4b, d and blue and red dots in Figs. 4c, e), indicating that measuring the velocity field ***v*** by PIV can effectively be replaced by the texture tensor method with high accuracy, consistently with the results in Sec. 4.3. More importantly, all three time series closely match each other, providing strong support for the cell number density equation (Eq. 8).

### 4.5 Evaluation of assignment rules of the strain rate decomposition

In the texture analysis, half-links that disappear and newly appear between two consecutive time frames are linked to specific morphogenetic cell events (Eq. 5; Fig. 1b). This assignment rule is not unique; for example, the approach proposed in Ref. [14] attributed the half-link from adjacent cells to a divided cell-to-cell division, it can also be assigned to a rearrangement (Supporting Information B and Fig. S5). To assess the impact of different assignment rules of the strain rate decomposition on the consistency of the time evolution equation for the cell number density Eq. 8, we conducted an evaluation (Supporting Information B for details). Our analysis revealed that alternative rules led to significant discrepancies with the cell number density equation, as shown in equation (Fig. S5). This suggests that the assignment rule proposed in Ref. [14] offered a consistent and optimal decomposition of the tissue strain to morphogenetic cell events.

### 4.6 Assessment of the kinematic equations

*Without topological deformation*. In the following analysis, we assess the kinematic relationship of **M**, represented in Eq. 7, where **M** is replaced by the experimentally measured 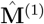. First, we examined the scenario without topological deformation (*i*.*e*., **D**_T_ = 0). In the absence of topological deformation, the change in texture tensor 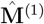 between consecutive time points obeys

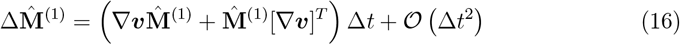

where Δ*t* denotes the time interval between frames (Eq. S13 in Supporting Information A). By utilizing data from ROIs with no topological changes in the connection of half-links (*that is*, no cell rearrangement, division, or apoptosis; 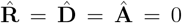), we evaluated the correspondence between the left- and right-hand sides of Eq. 16. Neglecting the higher-order terms of *𝒪* (Δ*t*^2^), we observed an excellent agreement (Fig. 5a and Fig. S6). The *𝒪* (Δ*t*^2^) term was sufficiently small, with a relative magnitude of approximately 1.2 *×* 10^−2^, considered negligible. These findings provide support for the kinematic equation, Eq. 7, in cases without topological deformation.

**Figure 5.**
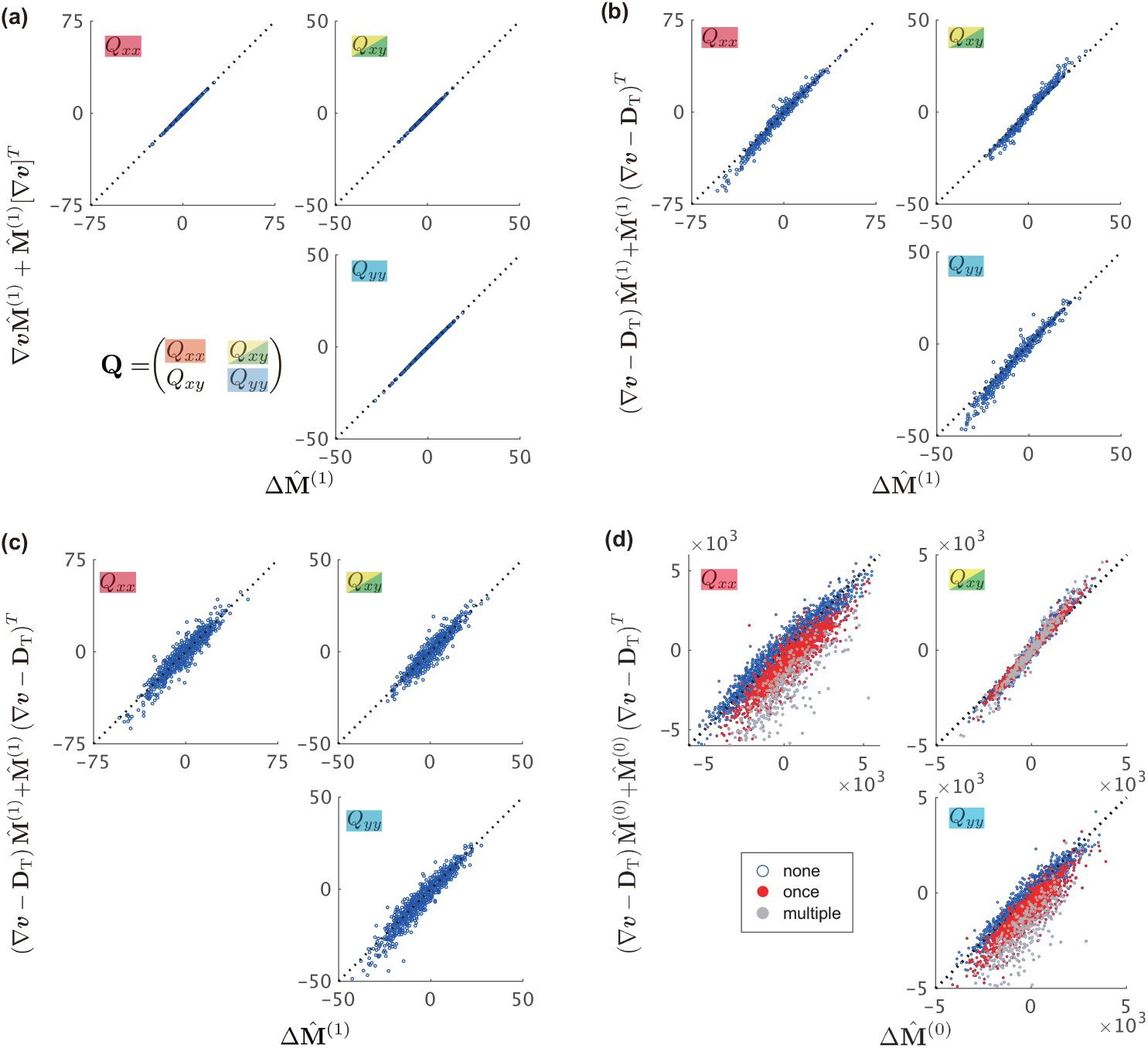
Accuracy of the kinematic equation for different definitions of 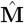. (a-d) The components of each tensor were evaluated for 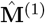 (a-c) and 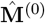 (d) using strain tensors based on 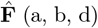 and PIV-measured ∇ ***v*** (c). Each point represents data from an individual ROI. In (a), only data from ROIs without topological deformations were plotted. In (d), point colors indicate the frequency of occurrence of cell division within the ROI: only once (red) or multiple times (gray) within the ROI.

The original definition of the texture tensor, **M**^(0)^, as well as the alternative definitions **M**^(2)^, **M**^(3)^, and **M**^(4)^ (Supporting Information C), yielded excellent agreement between the left- and right-hand sides of Eq. 16 (Fig. S7). This is because each term of ***l***_*ik*_ ⊗ ***l***_*ik*_ satisfied the kinematic equations and Eq. 7 remained unchanged up to a scaling factor of 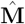.

*With topological deformation*. Next, we performed the same analysis using ROI data involving topological deformation. The kinematic equation for **M**, Eq. 7, was satisfied with minor deviations when the modified texture tensor 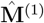 was employed (Fig. 5b, c and Fig. S8). This consistency was observed regardless of whether the strain rate tensors were determined using 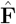 or PIV-measured ∇***v***.

In sharp contrast, the original texture tensor 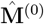 failed to satisfy the kinematic equation (Fig. 5d and Fig. S9). The degree of deviation from the identity line was notably influenced by the number of cell divisions (blue, red, and gray points in Fig. 5d indicate the data from ROIs with zero, one, and more than two divisions, respectively). This suggests that the deviation stemmed from a lack of proper normalization by the half-links *N*_h_ (Eq. 10).

Alternative definitions of the texture tensors 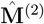 and 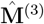 also resulted in larger deviations from the identity line compared with those obtained using 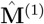 (Figs. S10a and b), which can be attributed to improper normalization. The appropriate normalization factor for representing the cell shape is the half-link number, *N*_h_, rather than the cell number, *N*_c_. 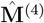 is in good agreement with Eq. 7 (Fig. S10c), comparable to the result obtained using 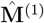. This is because the normalization factor ∼ *N*_c_ ∑ _*i*_ *n*_*i*_ in the definition of 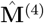 (Eq. S23) is of the same order as *N*_h_. These findings underscore the robustness of quantification when the texture tensor utilized for measurement is appropriately normalized. Overall, our data demonstrated that a properly normalized texture tensor, 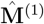, yielded quantitative data on tissue deformation that was highly consistent with the kinematic equations proposed in Ref. [19].

### 4.7 Reproducibility of the analysis among samples

To ensure the reproducibility of the aforementioned results, we replicated the measurements in two additional samples of the whole-wings and compared the outcomes across all three samples. First, we evaluated the consistency of results obtained using the texture tensor 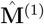. Kymographs of the cell shape characteristics during wing development in the region indicated in Fig. 6a are shown in Figs. 6b-d. The cell shape characteristics were derived from the texture tensor 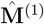, and we calculated the eigenvalues of 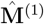, *λ*_1_, *λ*_2_, and indicated the cellular area 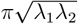 (Figs. 6b), as well as the cell aspect ratio 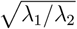 (Figs. 6c) and the local deviation in the direction of cell elongation (direction of the eigenvector for *λ*_1_) (Figs. 6d) [45]. These measurements exhibit spatial and temporal coherence across samples, validating the applicability of 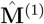.

**Figure 6.**
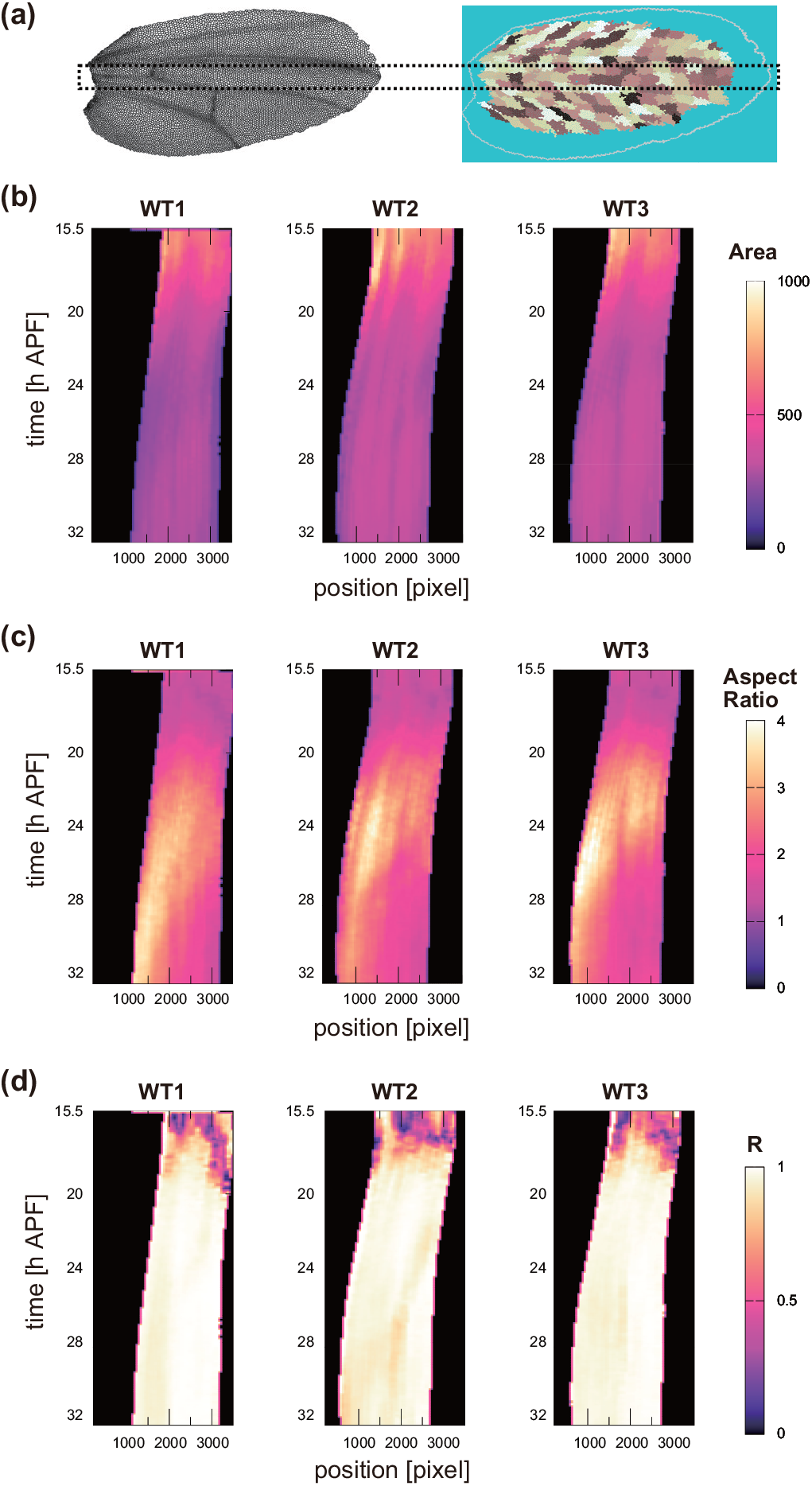
Reproducibility of cell shape quantities analyzed using the texture tensor across three wings samples. (a) Analyzed area of *Drosophila* wing (the area surrounded by the black dotted line). The area was set to include the C region. (b-d) Kymographs of cell shape quantities derived from the eigenvalues (*λ*_1_, *λ*_2_) and eigenvectors of the texture tensor 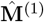. Quantities were calculated as moving average over a 200-px width (denoted by angle brackets) and are displayed at 50-px intervals. (b) Cell area, 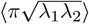. (c) Aspect ratio, 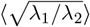. (d) Local deviation in direction of cell elongation, calculated by |⟨*e*^*i*2*θ*^⟩| where *θ* denotes the angle of the eigenvector associated with *λ*_1_.

Next, we validated the strain rates tensors derived from 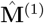. As previously mentioned, Figs. 3, S3, and S4 demonstrated consistency between the two ways of total strain rates measurements across different samples. Figs. 7a and b are times series of Tr**D**_T_ in the other samples, both of which are similar to Fig. 4b. Trace components of the quantified strain tensors 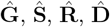, and 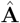 are also similar across different samples (Fig. 4a and Fig. 7c and d), supporting that the quantification procedure in our analysis is robust and reproducible.

**Figure 7.**
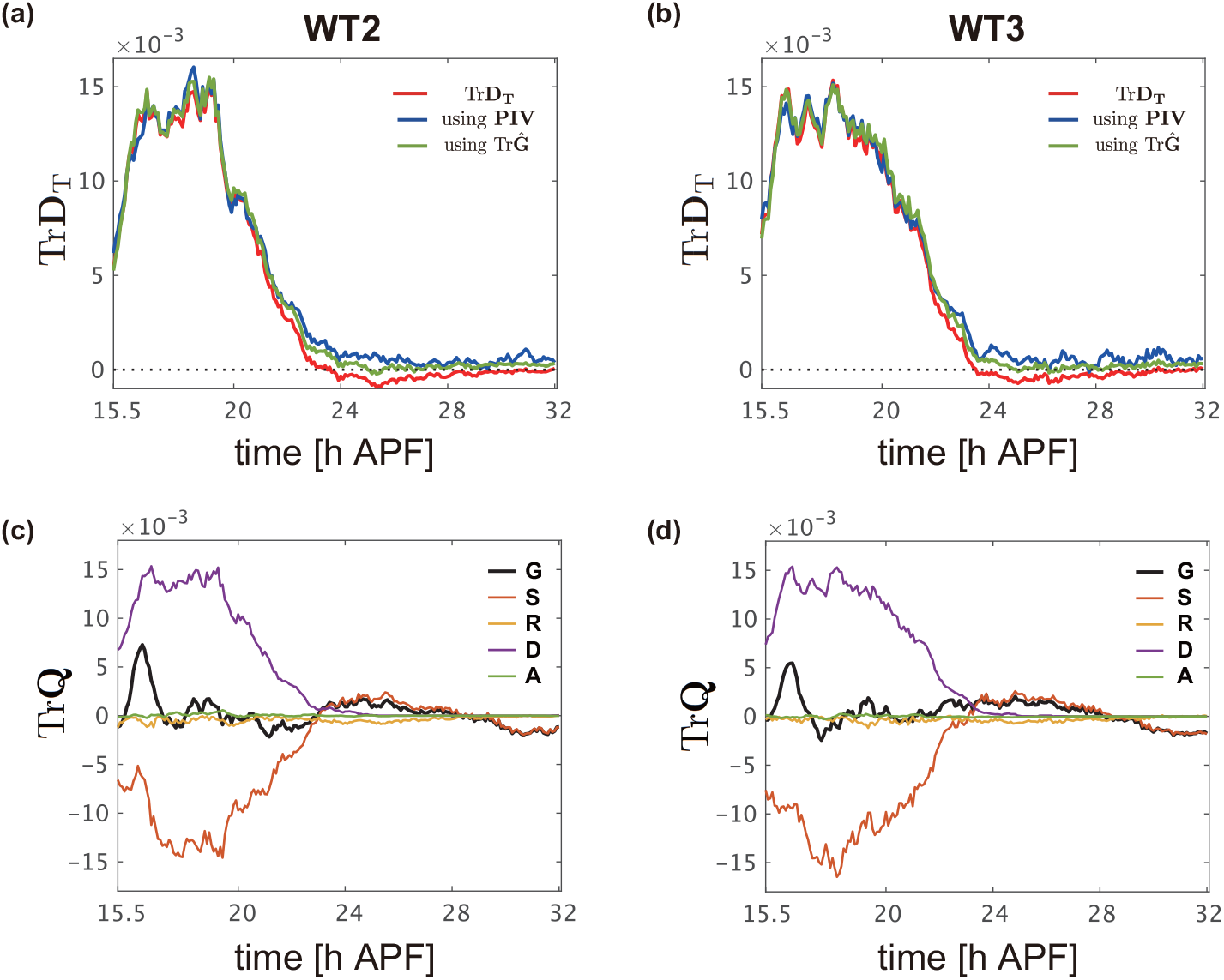
Reproducibility in validating the cell number density equations. (a, b) Time series data indicating the value obtained by dividing the left-hand side of Eq. 8 by 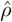 (blue and green lines), compared with Tr**D**_T_ (red line), for time-lapse data of the whole-wing captured at 5-min intervals starting from 15 h 30 min APF. (c, d) Time series of the traces of 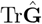 (black), Tr**Ŝ** (red), 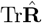 (yellow), 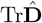 (purple), and 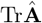 (green) for the same datasets as in (a, b). Data are from WT2 (a, c) and WT3 (b, d). Sample IDs are similar to those in Fig. 6.

## 5 Discussion

Kinematic relationships play a key role in the dynamics of deforming materials, as they typically involve only a few parameters and represent the general and robust characteristics of a material compared to kinetics. In this study, we assessed the kinematic equations governing the cell shape tensor (Eq. 7) and cell number density (Eq. 8) using experimental data from *Drosophila* wings. To accomplish this, we introduced several potential definitions for the texture tensor, among which **M**^(1)^ demonstrated a good agreement with direct pixel counting for cell area and with the second moment for cell shape. We also validated that the total strain tensor 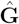 , derived from the change in **M**^(1)^ between consecutive time frames, properly aligned with that obtained through PIV. By leveraging **M**^(1)^, we demonstrated the compatibility of the kinematic equations proposed in Ref. [19] with experimental data, thereby validating the theoretical framework of the kinematic equations while also demonstrating the practical utility of the texture tensor **M**^(1)^ as a reliable indicator for analyzing morphogenetic cell events.

Our analysis revealed that all definitions of the texture tensors, **M**^(*m*)^ (*m* = 0, 1, 2, 3, 4), support the kinematic equation for **M** (Eq. 7) in the absence of topological changes in the texture (Fig. 5a and Fig. S7). However, when topological changes in the texture were involved, only the analyses using **M**^(1)^ and **M**^(4)^ successfully supported the kinematic equation, whereas those using other definitions, including the original **M**^(0)^, did not. This discrepancy is attributed to the selection of normalization factors. In the presence of topological changes, the number of half-links changes, necessitating an update to the normalization factors. The analysis using **M**^(4)^ yielded results comparable with those using **M**^(1)^ (Fig. S2d and Fig. S10c) owing to their normalization factors being of similar magnitude (*N*_h_ ∼ *N*_c_ ∑ _*i*_ *n*_*i*_), rendering them invariant to the size of the ROI. This finding further emphasizes that the specific definition of the texture tensor is not crucial as long as an appropriate predetermined normalization factor is applied, hence underscoring the robustness of the analysis using the texture tensor.

The kinematic equation for **M**, Eq. 7, was derived assuming the epithelial tissue can be modeled as a tiling of polygons, where changes in cell shape are tightly coupled with tissue strains [18, 19]. In this equation, cell shape changes are assumed to be governed solely by tissue strains, ∇***v***, and topological changes, **D**_T_. For example, in the absence of topological changes (**D**_T_ = 0), cell shape changes 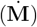 and total tissue strain (∇***v***) must remain compatible with each other. Note that this kinetic relationship differs from that of other systems such as liquid crystals [46], in which molecular alignment, represented by the nematic tensor, can evolve independently of the macroscopic velocity, as indicated by appearance of a relaxation term in the time evolution equation for the nematic tensor. Furthermore, the cell number density equation Eq. 8 is directly derived from the kinematic equation Eq. 7 as discussed in Sec. 2.2. Our data analysis using the revised texture tensor method confirmed that both equations (Sec. 4.4 and Sec. 4.6) are highly consistent with quantitative data of tissue deformation and morphogenetic cell events.

The current texture tensor analysis procedure for decomposing strain rates, as described in [14] and Supporting Information A, has room for further improvement. Specifically, the method does not account for rotational deformation, and the choice of normalization in deriving Eq. S14, as well as the definition of the residual term Ψ in Eq. S17, lack direct interpretability in terms of cellular or continuum deformation kinematics. While a deeper exploration of the theoretical framework lies beyond the scope of the present study, it represents an important and intriguing avenue for future research. We also note that in situations where cell-cell connections are weaker or looser, such as in mesenchymal tissues, modifications to both the kinematic equations and the corresponding strain measurement methods may be required.

Having established the kinematics, kinetic relationship should be addressed. However, validating kinetics requires more advanced techniques, as it involves numerous parameters in constitutive equations that, unlike kinematics addressed in this study, may not be directly measurable. One promising approach is data assimilation, a statistical technique that integrates computational model with observed data [47]. By iteratively combining simulation and fitting, data assimilation enables not only quantitative, data-driven simulation but also system identification, including the evaluation of model parameters, their uncertainties, and the goodness-of-fit of the model [48, 49]. This method has been successfully applied to biological systems [50–53]. The precision in the kinematic relationships achieved through our texture tensor analysis provides a solid foundation for such an approach. However, implementing and testing it requires substantial effort and will be the focus of future studies.

In conclusion, we cross-validated the kinematic equations of multi-scale continuum model and the strain measurement methods for epithelial tissue. The approach used in this study can be useful for testing other related methods that employ different kinematic and kinetic frameworks. We anticipate that the quantitative comparison and validation of our methods, in conjunction with related approaches, will aid in bridging tissue- and cell-scales in epithelial morphogenesis.

## Author contributions

T.N., K.S. and S.I. designed the research. T.N. and S.I. performed reformulation of the tensor analysis. K.S., S.I., and T.N. conducted the image segmentation. T.N. performed the image and data analyses. K.S. performed the experiments. T.N., K.S., and S.I. drafted the manuscript. All authors approved the final manuscript.

## Data availability

The data are available from the lead contact upon reasonable request. The code for texture analysis can be downloaded from https://github.com/IshiharaLab/TextureTensorAnalysis.

## Acknowledgments

The authors thank Yang Hong for providing reagents; Kyoko Komano and Miho Aruga for their assistance with image segmentation; and Philippe Marcq, François Graner, Yohanns Bellaïche, and Boris Guirao for their valuable discussions and critical review of the manuscript. This study was financially supported by JST CREST (JPMJCR1923), JPJSBP (JPJSJPR 201915010), and JSPS Kakenhi (24H01931) to S.I., and the AMED PRIME program (20gm5810025h9904) and JSPS Kakenhi (24K02029 and 25H01364) to K.S.

## Supporting Information

### A Tissue deformation analysis by texture tensors

We summarize the texture tensor analysis utilized in the main text to quantify cell and tissue deformations. This method involves calculating strain tensors that result from morphogenetic cell events by examining temporal changes in the texture tensor 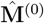 (Eq. 10 of the main text). While our approach is based on the method outlined by Guirao et al. [1], we offer an alternative derivation of the deformation gradient tensor, 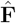 (Sec. A2). Furthermore, the specific expression of the strains to be measured differs slightly from the previous ones, with the deviation being of the order *𝒪* (Δ*t*^2^) (Sec. A3).

The data analysis workflow is summarized in Fig. S1. In this study, coarse-grained measurement were performed using ROIs defined by cell-tracking data (*i*.*e*., the same cells are tracked from the initial to the final time points; Sec. 3.7 and Sec. 3.8 in the main text). This allows for evaluation of temporal changes in the cell shape field 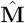 without accounting for influx and efflux, in other words, we evaluate the Lagrange derivative of 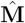 at each time point.

#### A1 Temporal changes in the texture tensor

The change in 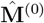 between two consecutive time frames is defined as follows:

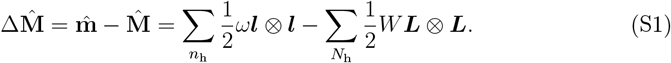

In this expression, uppercase and lowercase letters represent quantities measured at the earlier and later time points of the consecutive frames, respectively (*i*.*e*., at time *t* and *t*+Δ*t*). With the cell-tracking data, 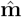 and 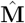 are calculated from ROIs composed of the same cells, or their mother or daughter cells (Sec. 3.7 in the main text; Fig. S1c, d). Thus, 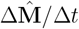 evaluates Lagrange derivatives 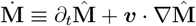 at time point *t* (Sec. 3.8). At time *t* + Δ*t*, the total number of half-links *n*_h_ is the sum of the number of conserved links, *n*_c_, and the number of links that appeared, *n*_a_, between the time frames. Similarly, at time *t*, the total number of half-links *N*_h_ is the sum of the number of conserved links, *N*_c_ = *n*_c_, and the number of links that disappeared, *N*_d_. The decomposition of Eq. S1 is as follows:

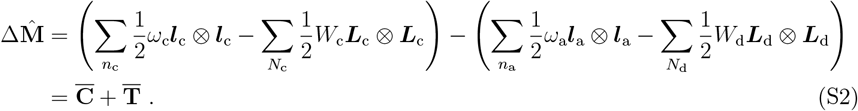

The first term enclosed in brackets in Eq. S2 comprises links that maintain their neighboring relationships, denoted as 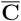. The second term, 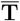, signifies the overall change attributed to the topological processes and can be decomposed as 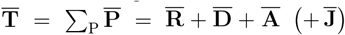 , where 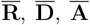, and 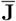 indicate that rearrangement, division, apoptosis, and flux (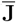appeared only in the Eulerian description and is absent in our analysis). The abbreviations 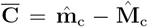 and 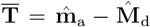 are utilized. These tensors have squared length dimensions.

The decomposition of 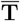 into 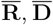 and 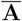 was performed as follows: 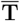 was calculated from the links that disappeared or appeared between consecutive frames. As explained in the main text, each link comprises two half-links ***l***_*ik*_ and ***l***_*ki*_(= −***l***_*ik*_) (or ***L***_*ik*_ and ***L***_*ki*_ = −***L***_*ik*_), with both half-links belonging to the same link being assigned to the same morphogenetic cell event. The allocation of half-links to division and apoptosis took precedence over rearrangements. An example of the assignment is shown in Fig. 1b. For further discussion on this topic, refer to Sec. 4.5 in the main text and Sec. B in the Supporting Information.

#### A2 Deformation gradient tensor F for tissue deformation

Deformation of a continuum material can be described using a deformation gradient tensor **F** [2]. In our texture tensor analysis, we calculate the empirical deformation gradient tensor 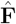 based on the half-links obtained through *in vivo* measurements. Consider the deformation of a continuum object, where a material point ***r*** at time *t* is mapped to ***R*** = ***r*** + ***u*** at time *t* + Δ*t*, with ***u*** representing the displacement vector. The relative position *d****r*** between two points at infinitesimal distances changes to *d****R*** = **Fd*r*** owing to deformation, where **F** is the deformation gradient tensor defined as

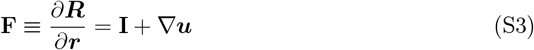

with the identity matrix **I**.

In practice, **F** is evaluated using half-links contained in the corresponding ROI between consecutive time frames. When the tissue deforms without topological changes 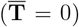, all links are conserved between two consecutive time frames. In such cases, a conserved link changes from ***L***_c_ to ***l***_c_, satisfying ***l***_c_ = **F*L***_c_. Even in cases involving topological changes 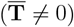, we can assume that most links are conserved, with only a small fraction of half-links appearing and disappearing. Assuming affine deformation in each ROI and a constant **F, F** is determined from the experimental data by minimizing the function

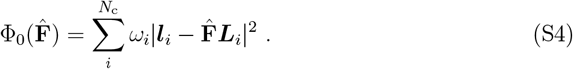

where the summation is performed over the conserved links. **F** is estimated as follows:

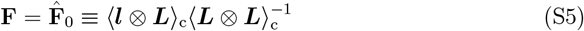

where 2*×*2 tensor ⟨***l*** ⊗ ***L***⟩_c_ is defined by 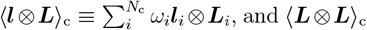, and ⟨***L*** ⊗ ***L****⟩*_c_ is defined similarly. Furthermore, **F** can be determined by considering an alternative function^1^;

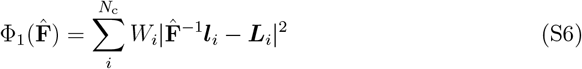

and 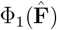 is minimal at

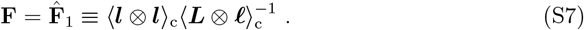

Notably, 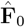 and 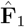 satisfy the following relationships:

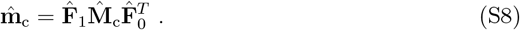

Furthermore, because 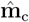 and 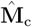 are both symmetric tensors, *that is*, 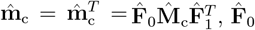 and 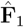 satisfy 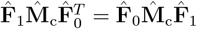.

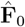and 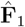 represent two empirical approximations of the deformation gradient tensor **F**, anticipated to have a similar construction. Moreover, they should align in the case of the ideal pure affine deformation, resulting in the following relationship:

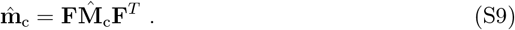

We chose the deformation gradient tensor as the arithmetic mean in our analysis.

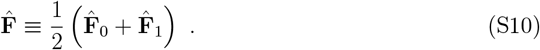

This quantity, calculated from conserved half-links in the ROI, was utilized for the analysis discussed in the main text.

We evaluated the relative mismatch using data from *Drosophila* epithelial tissues (pupal wing and notum).

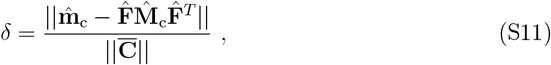

where the norm of the second-order tensor **a** is defined as 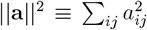. The mismatch *δ* is sufficiently small (*<* 4.0 *×* 10^−3^), validating the appropriateness of 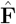 as the definition of the deformation gradient tensor. Moreover, we examined the mismatch using the geometric means of 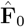 and 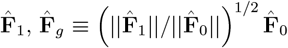 , instead of 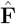. The small mismatch (*<* 8.0 *×* 10^−3^) between 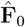 and 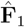 suggests that they are similar, and our analysis results are not significantly influenced by the choice of their means. The symmetric part of the total strain-rate tensor is calculated using 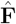 as follows:

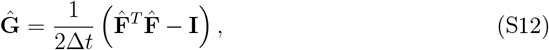

which was utilized in our analysis (Eqs. 2, 4, and 11 in the main text).

For deformations without topological processes, 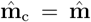 and 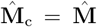 applies. Eqs. S3 and S9 lead to 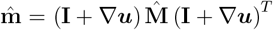 , and then 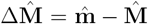 (Eq. S1) reads

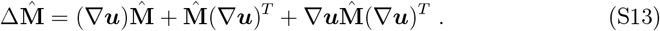

The equation obtained by omitting the third term on the right-hand side (higher-order term with respect to Δ*t*) corresponds to Eq. 16 in the main text.

#### A3 Dimensionless symmetric strain rate tensors

The strain rate tensors **G, S, R, D**, and **A** utilized in the continuum theory have units of the inverse of time. We outline the calculation of the corresponding empirical strain tensors 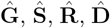, and 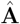 from experimental data. Assuming the preservation of most links, the conserved half-links are utilized to derive the deformation gradient with 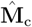 as the reference state. The dimensionless symmetric tensors for 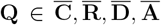 are defined as follows:

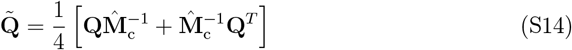

where the tilde denotes an operation that produces dimensionless symmetric tensors using 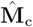. 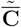 is calculated as

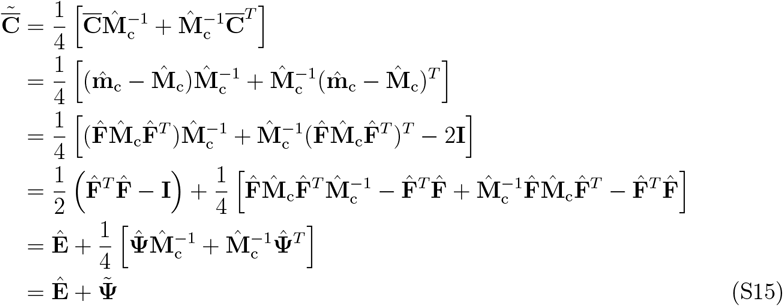

where we utilized

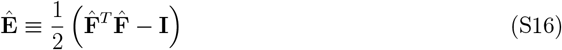

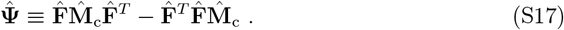

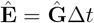 represents the Green-Lagrange strain tensor with respect to the deformation between two consecutive time frames [4]. 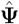 is expressed as 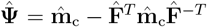, and vanishes if 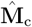 and 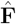 commute.

Ref. [1] adopted 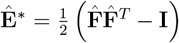 as a measure of deformation instead of 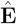. The difference, 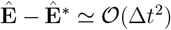, is negligible (Fig. S11).

#### A4 Decomposition of the strain rate to cell morphogenetic events

The tissue strain tensor 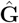 defined in Eq. S12 can be decomposed into the strains res-ulting from cell morphogenetic events. By substituting Eq. S15 into Eq. S2, we obtain

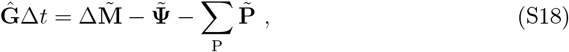

where Eq. S14 was employed. Here, 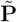 represents the contribution from topological events, with each 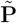 calculated from the half-links assigned to the corresponding topological cellular event. 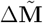 represents the total deformation of the ROI in terms of the size and shape, which is further partitioned into components that correspond to the respective cellular events. Notably, *N*_h_ = *N*_c_ + *N*_d_ and *n*_h_ = *n*_c_ + *n*_a_ represent the numbers of half-links at time *t* and *t* + Δ*t*, respectively. *N*_d_ and *n*_a_ denote the numbers of disappearing and appearing half-links associated with topological cellular events, respectively. These are further decomposed into *N*_d_ = ∑_P_ *N*_P_ and *n*_a_ = ∑_P_ *n*_P_, where the subscript P denotes either T, D, or A. The change in the number of half-links expressed as Δ*N* = *n*_h_ − *N*_h_ = *n*_a_ − *N*_d_ = ∑_P_ (*n*_P_ − *N*_P_) ≡ ∑_P_ Δ*N*_P_. The deformation is of the ROI from 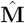to 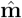 is partitioned based on the numbers of half-links, as follows:

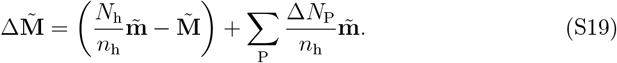

The magnitude of 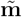 is normalized by *N*_h_*/n*_h_, rendering it comparable with 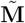 . The residual fraction of 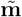 in the last term is attributed to topological cellular processes. Notably, the value of Δ*N*_P_ is expected to be positive for division (D), negative for apoptosis (A), and Δ*N*_P_ *≃* 0 for rearrangement (R) (Fig. 4 for the experimental validation).

These arguments enable the decomposition of tissue deformation into contributions from each cellular event. From Eqs. S18 and S19, we obtain:

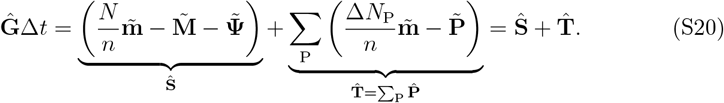

The notation **D**_T_ utilized in the continuum model [5] corresponds to 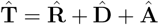.

### B Inconsistencies in cell number density equations resulting from the inappropriate assignment rules

In the analysis of texture tensor, each half-link is associated with a specific cell morphogenetic event: cell shape change (S), rearrangement (R), division (D), or apoptosis (A) (Fig. 1a). However, determining the assignment rule can be complex, particularly when dealing with topological changes in half-link connections during R, D, and A events. Consider a scenario in which a cell divides, as shown in Figs. S5a. The dividing cell(s) and their first-neighbor non-dividing cells are colored in green and gray, respectively. Cells are categorized based on changes in their relationships with neighboring cells. The centers of dividing cells are denoted by light green points. For first-neighbor non-dividing cells, the centers of cells with an increased number of adjacent cells are represented by blue points, whereas those without such changes are denoted by black points. The question arises: to which cell morphogenetic event should the half-link between dividing and non-dividing cells be assigned? In ref. [1] and the main text of this study, the half-links are considered undirected edges, with both half-links between the pairs of cells attributed to the same morphogenetic event. Alternatively, considering half-links as directed edges could lead to a rule dependent on direction.

We assessed whether the consistency of the time evolution equation for cell number density (Eq. 8 in the main text) was influenced by the different assignment rules of the strain-rate decomposition. The assignment rule for the half-links adopted in the main text is shown in Fig. S5b. We reproduced the results shown in Fig. 4b. The results of the time evolution in Eq. 8 are shown in Fig. S5c with direction-dependent assignment rules applied. These rules include: (i) Assigning all half-links from white-dot cells to cell division. (ii) Assigning half-links from blue to white-dot cells to cell division owing to the division of the opposing cell. (iii) Considering half-links from black to white-dot cells as cell shape changes reflecting the consistent relationship with the opposite cell. (iv) Classifying half-links between black-dot and blue-dot cells as cell shape changes. The application of these rules resulted in a time series of cell number density that did not align with those obtained by substituting the deformation field with data from PIV.

Furthermore, we explored a scenario in which the assignment is independent of the direction of the half-links; however, the rules differ from those utilized by Guirao et al. [1]. The following assignment rules were employed: (i) Links between (white, white) and (white, blue)cells were assigned to cell division. (ii) Links between (black, white) and (black, blue)cells were assigned to cell shape changes. As shown in Fig. S5d, these rules resulted in a greater discrepancy compared with that observed in Fig. S5c, likely resulting from the underestimation of strain from topological deformation.

### C Alternative definitions of the texture tensor

The texture tensor was introduced in the form of Eq. 10 and is modified as Eq. 13 in the main text. We also considered other possible forms of the texture tensor:

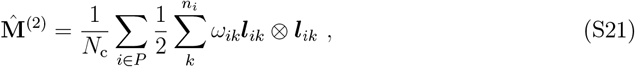

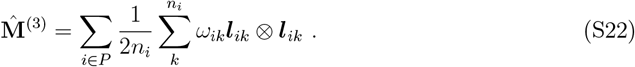

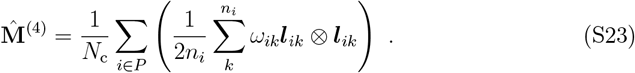

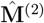 is normalized by the number of cells in the ROI. 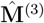 and 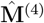 take into account the polygonal class of cells using a weighting factor proportional to 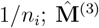 is normalized by neighboring cells *n*_*i*_ to equalize the contribution of each polygonal cell. 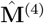 is further normalized by the cell number in the ROI, *N*_c_, and is interpreted as a mean of individual cellular shape tensor 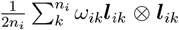 over the ROI. All the proposed definitions of texture tensors possess a physical dimension of squared length, differing primarily in the normalization procedure based on the number of cells and their adjacent counterparts.

---

**Figure S1.**
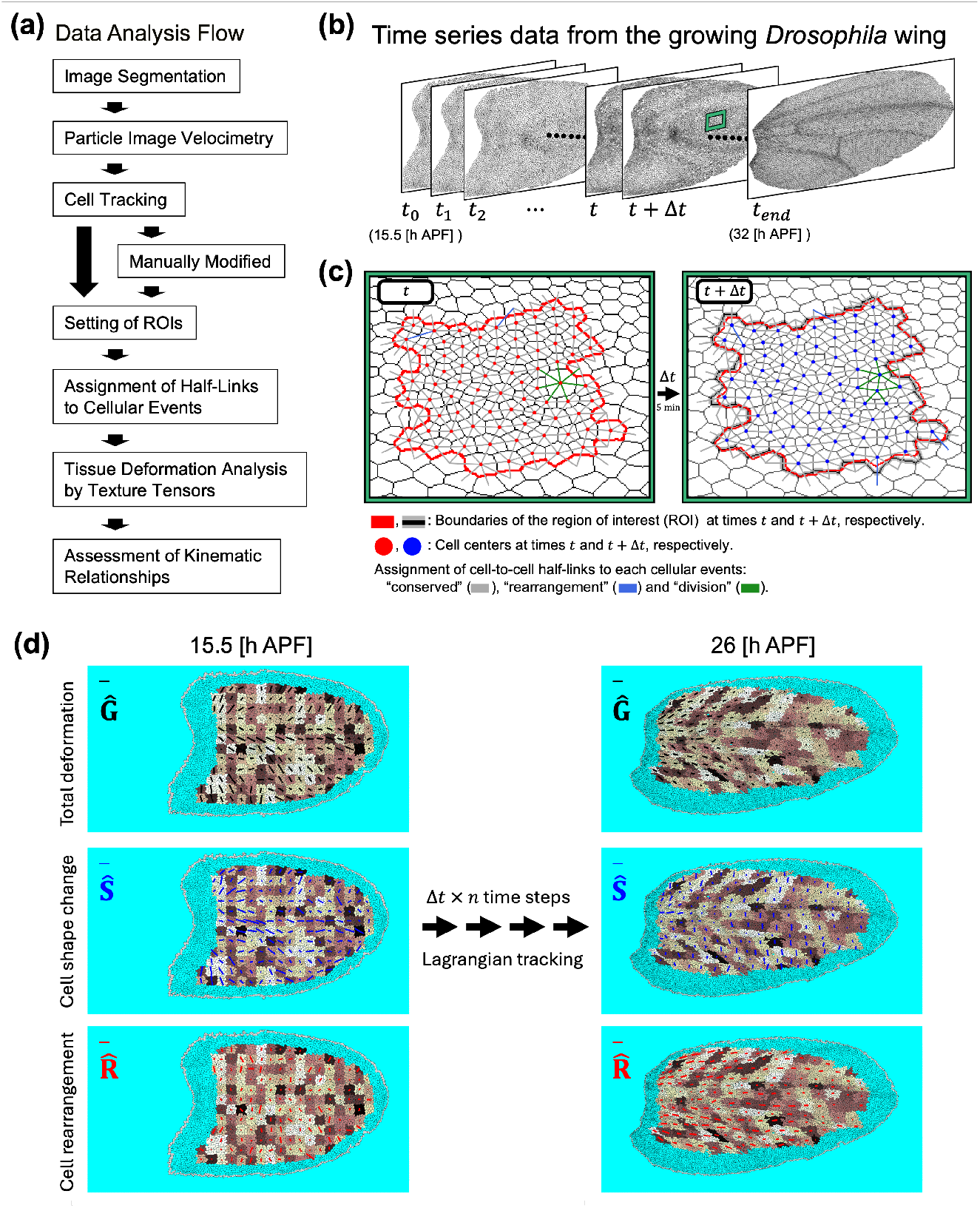
Schematic diagram of data analysis. (a) Flowchart of the data analysis procedure. (b) Skeletonized time-series image of growing *Drosophila* wings. (c) Changes in the texture in the region of interest (ROI) indicated by the closed red lines. Half-links between cell centers at times *t* and *t* + Δ*t* (red and blue filled circles, respectively) are shown. Color of half-links indicate their assignment to morphogenetic cell events (gray: conserved, blue: rearrangement, and green: division). The gray line represents the ROI boundary at each time point, while the red line at time *t* +Δ*t* is used for comparison with the ROI bounrady at *t*. (d) Spatial maps of mean-field quantities representing total deformation (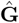, black lines), cell shape change (**Ŝ**, blue lines), and cell rearrangement (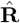, red lines) across the entire wing at 15.5 and 26 h APF, with 15.5 h APF taken as the initial time. Bar shown in each ROI represents the deformation rates of the respective cellular events derived from the deviatoric component of each strain rate. The reference line in the top-left corner of each panel corresponds to a 1% change over 5 minutes. Time averaging was performed over 2-hour intervals.

**Figure S2.**
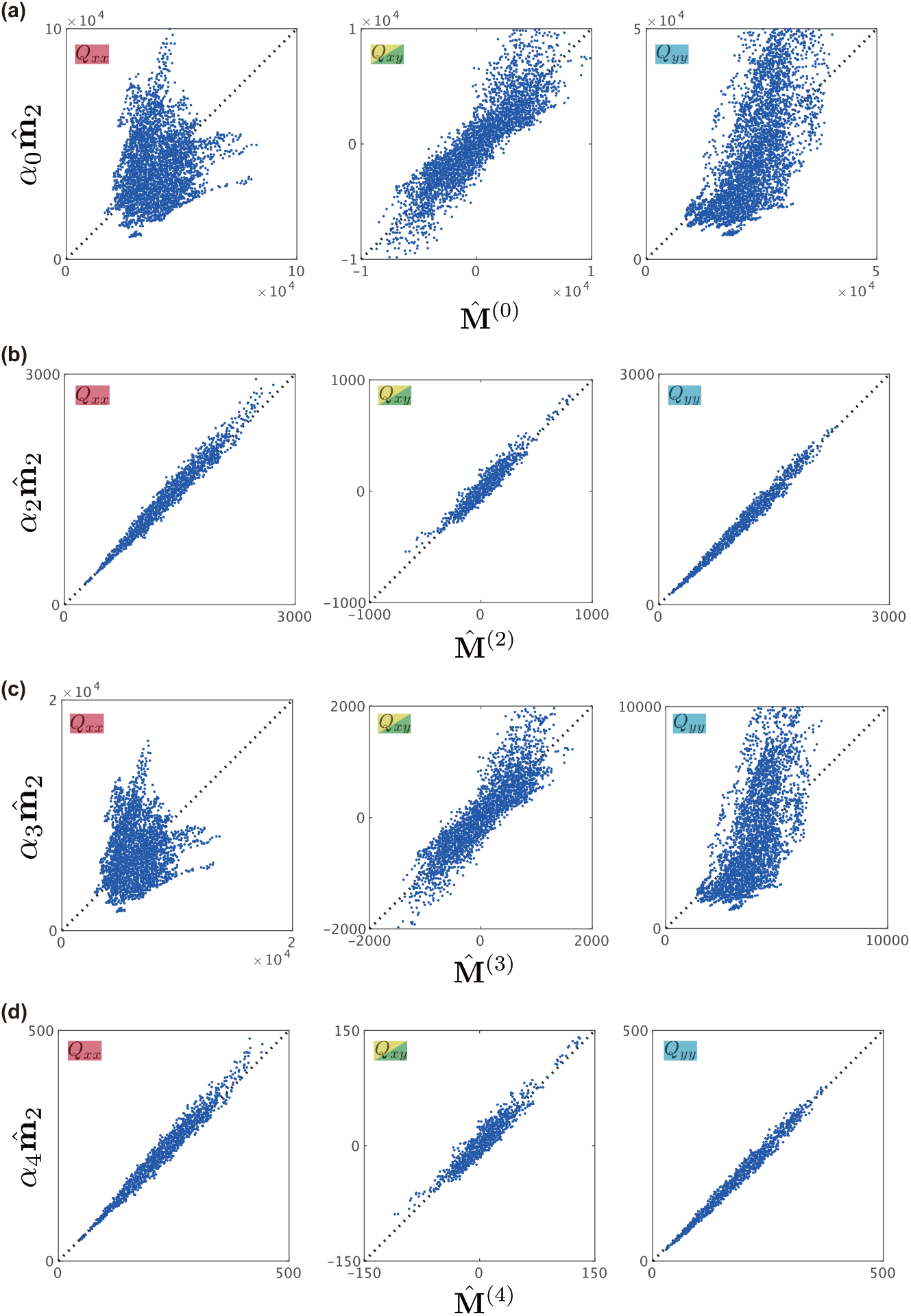
Comparison of the second moments of cell shape for different definitions of cell shape tensors, 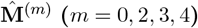. (a–d) Each component of the tensors is shown in the respective panels. Individual dots represent data from single ROIs of 120 pixels × 120 pixels from the entire wing images (hereafter, the average is obtained over ROIs of this size unless noted otherwise). A scaling factor *α* was introduced as a fitting parameter for each measurement: (a) 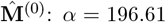, (b) 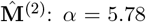, (c) 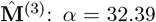, and (d)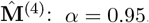 .

**Figure S3.**
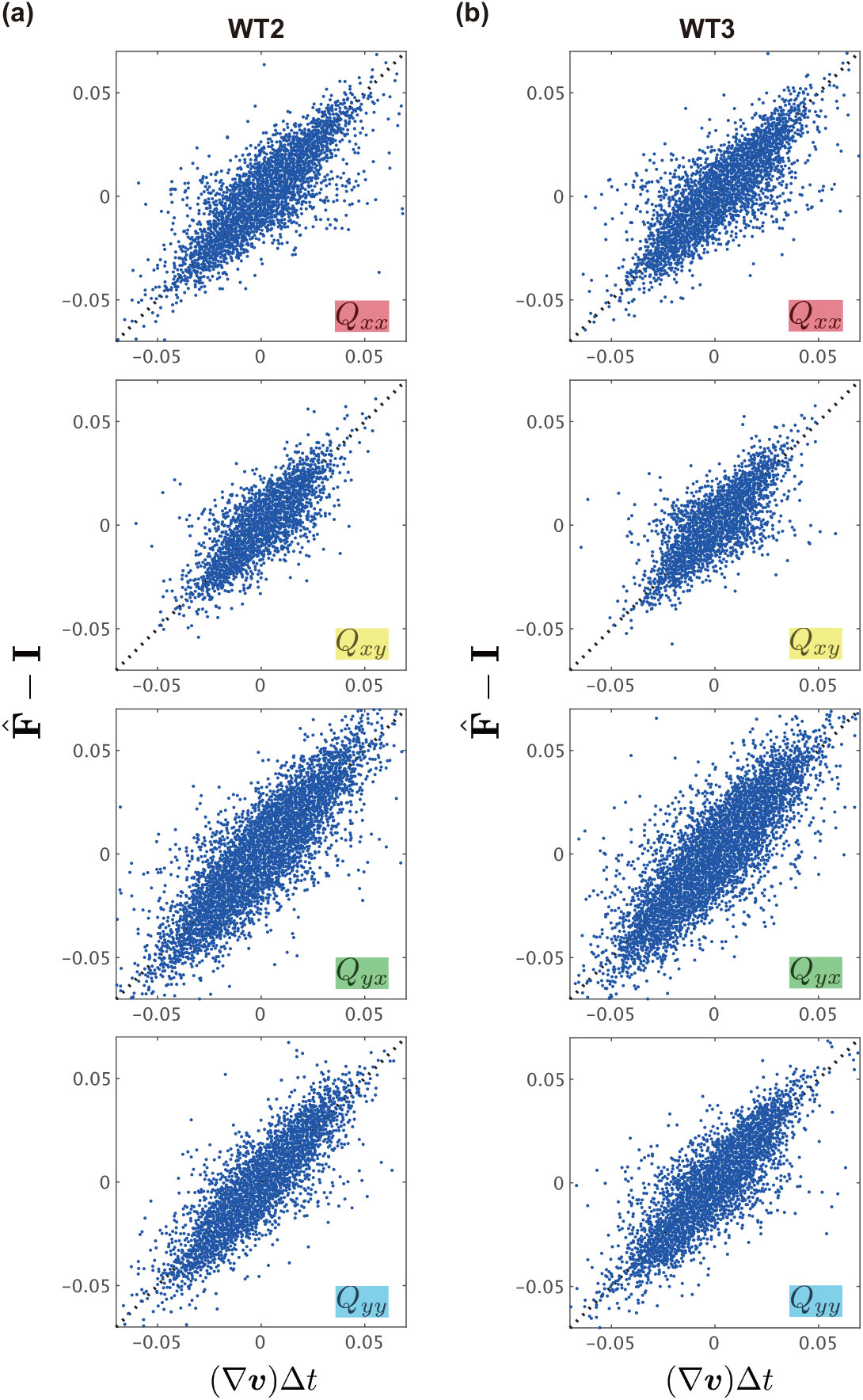
Validation of strain rate tensors using data from additional samples. (a, b) Data from WT2 (a) and WT3 (b) are analyzed and plotted similarly as in Fig. 3a.

**Figure S4.**
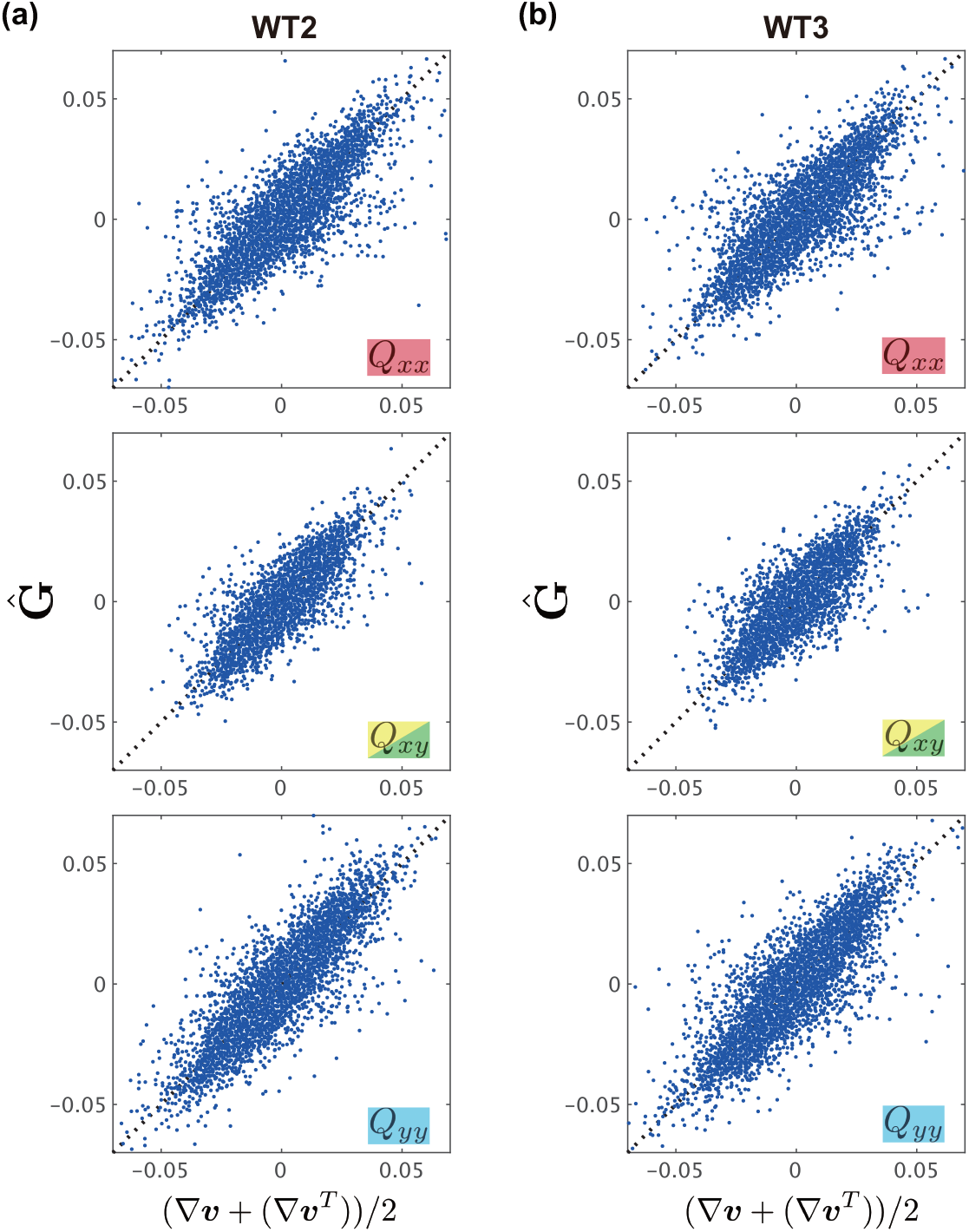
Validation of the symmetric part of strain rate tensors using data from additional samples. (a, b) Data from WT2 (a) and WT3 (b) are analyzed and plotted similarly as in Fig. 3b.

**Figure S5.**
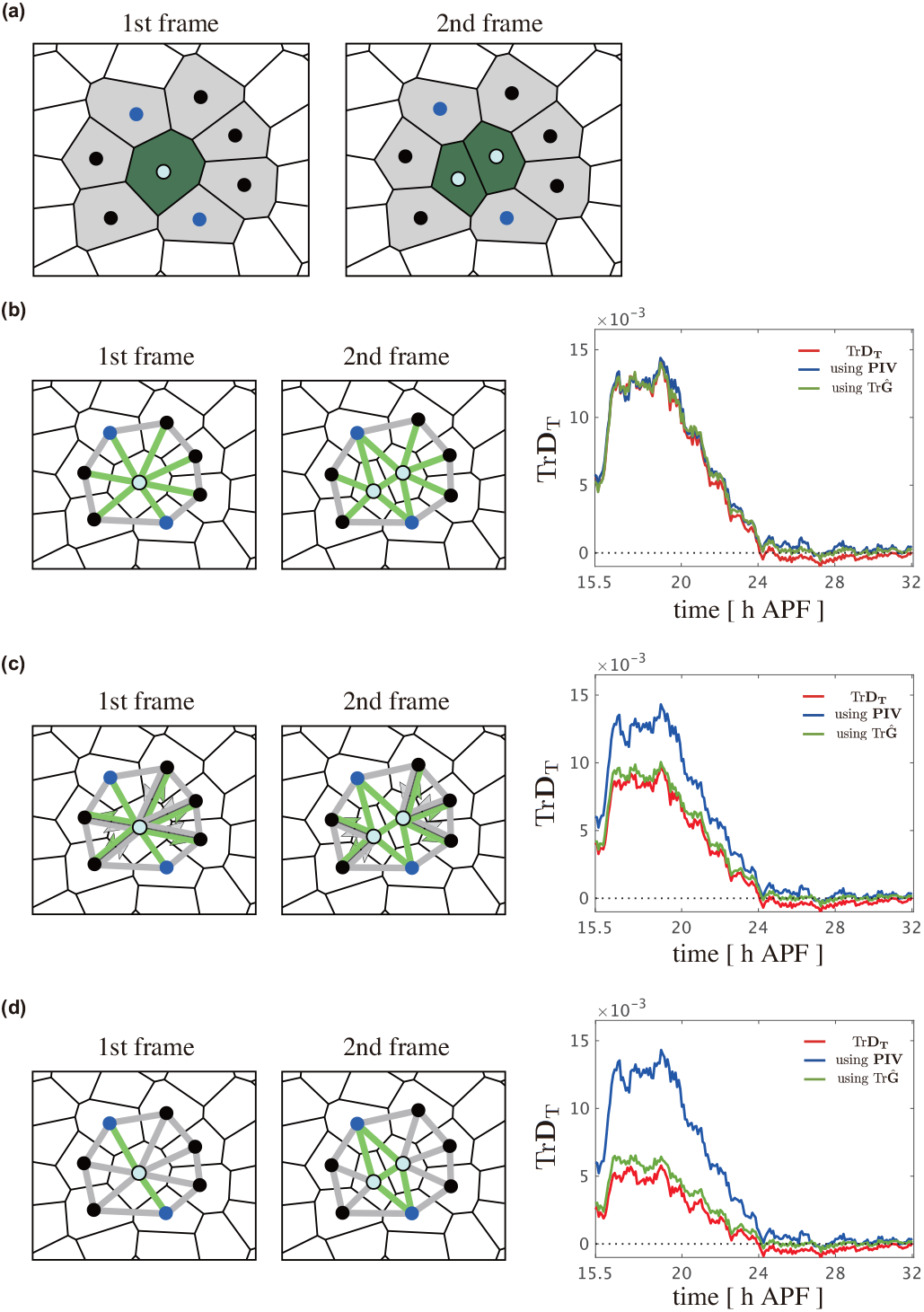
Evaluation of assignment rules of half-links to each cellular event. (a) Illustration of cell geometry change resulting from a cell division between the first to second timeframes. The dividing cells and their first-neighbor, non-dividing cells are distinguished by their colors (green and gray, respectively). The centers of the dividing cells are indicated by light blue points. In the case of first-neighbor, non-dividing cells, those with an increase in the number of adjacent cells are indicated with blue points, whereas those without such changes are represented by black points. (b-d) Tests of the cell number density equation (Eq. 8 in the main text) using different assignment rules. The left panels illustrate the assignment rules for half-links, with gray and green lines indicating half-links assigned to “conserved” and “division,”. (b) Rule employed in the main text (the right panel is identical to Fig. 4b in the main text). (c, d) Alternative assignment rules for half-links involved in division (Sec. B). In and (d), undirected links are assigned to the same cellular event for both directed half-links. In (c), half-links indicated by bi-directed arrows are assigned to different cellular events depending on their direction. The right panels indicate time-series data obtained by utilizing the corresponding assignment rules. The same whole-wing data (WT1) as in Fig. 4 was utilized, with data plotted similarly as in Fig. 4b. The values obtained by dividing the left-hand side of Eq. 8 by 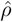 are represented by blue and green lines and are compared with Tr**D**_T_, represented by the red line.

**Figure S6.**
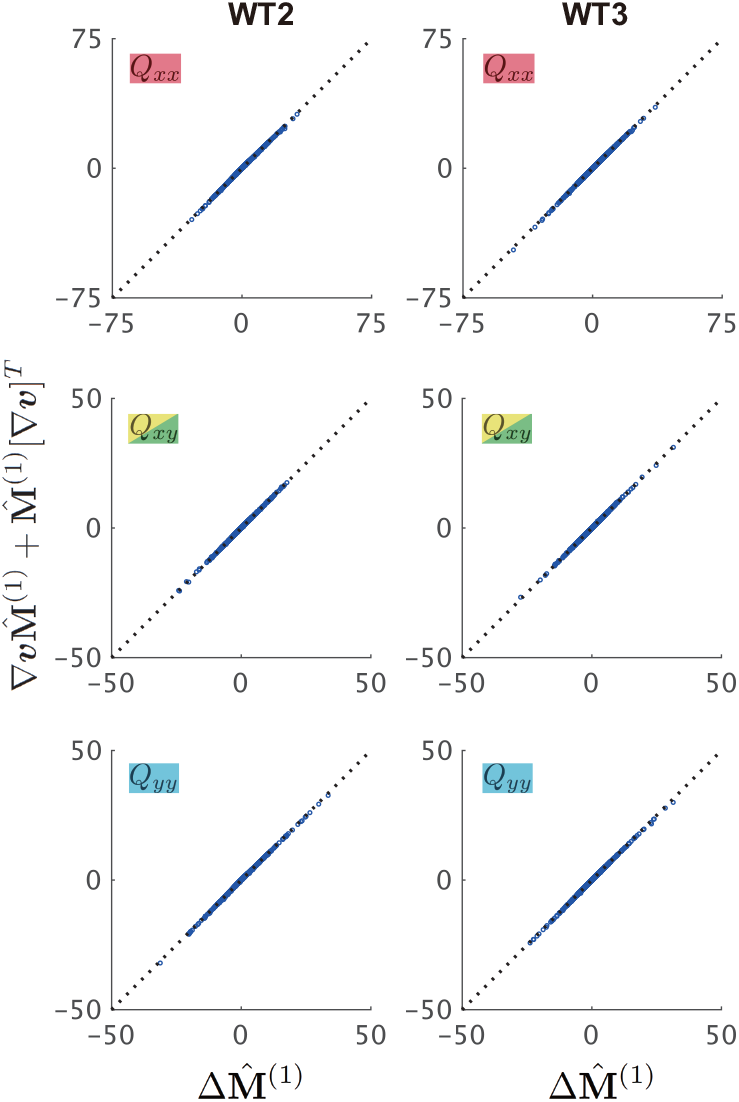
Additional data for the validation of the kinematic equation 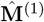 in ROIs without topological deformation. Data from WT2 and WT3 wings (left and right columns) are analyzed and plotted similarly as in Fig. 5a.

**Figure S7.**
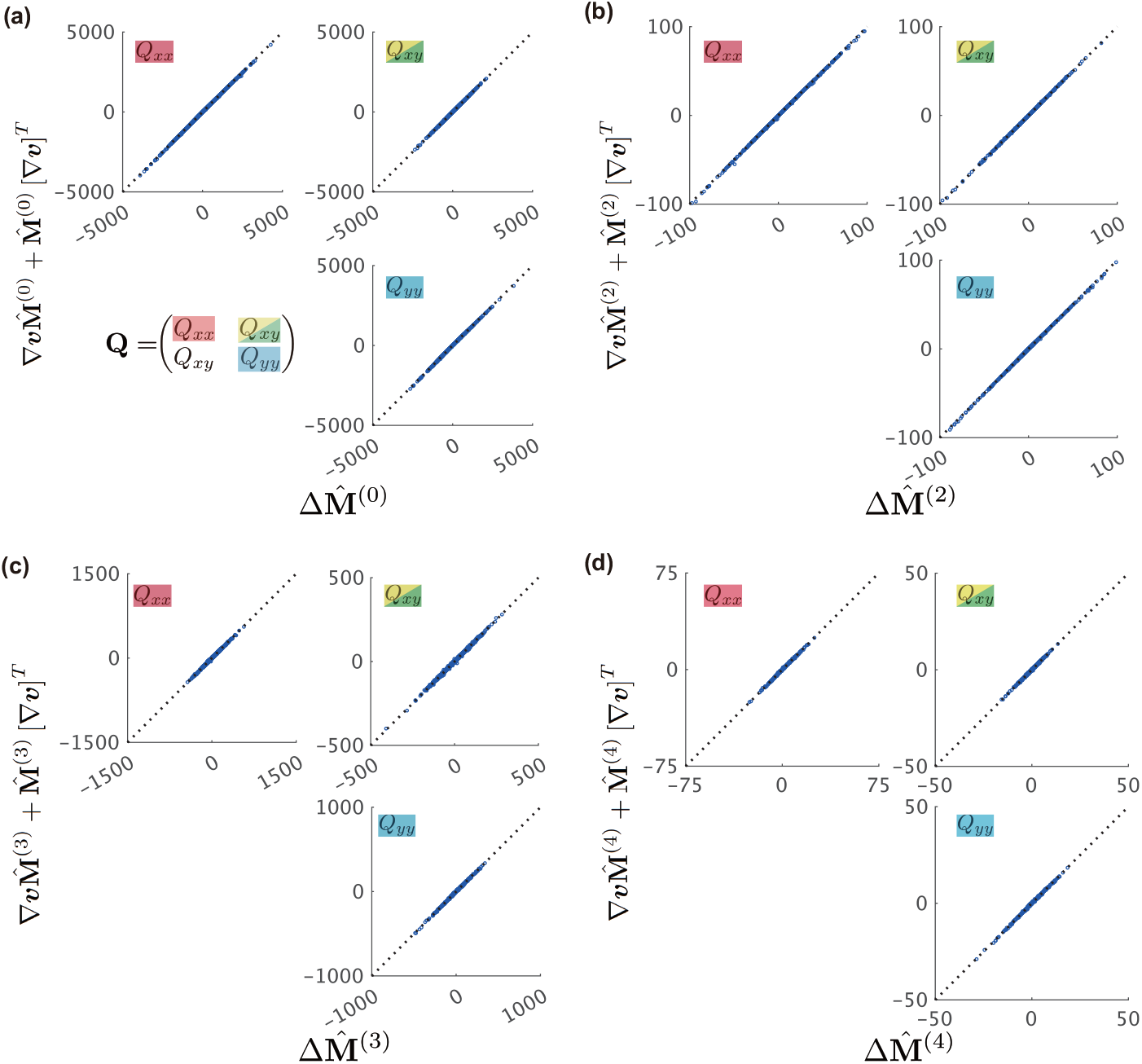
Validation of kinematics for various definitions of 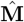 in ROIs without topological deformation. (a–d) The components of each tensor were evaluated for the following definitions of the texture tensor: (a)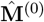, (b)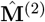, (c)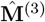, and (d)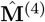. The same whole-wing data (WT1) as in Fig. 5a was utilized, with data plotted similarly as in Fig. 5a.

**Figure S8.**
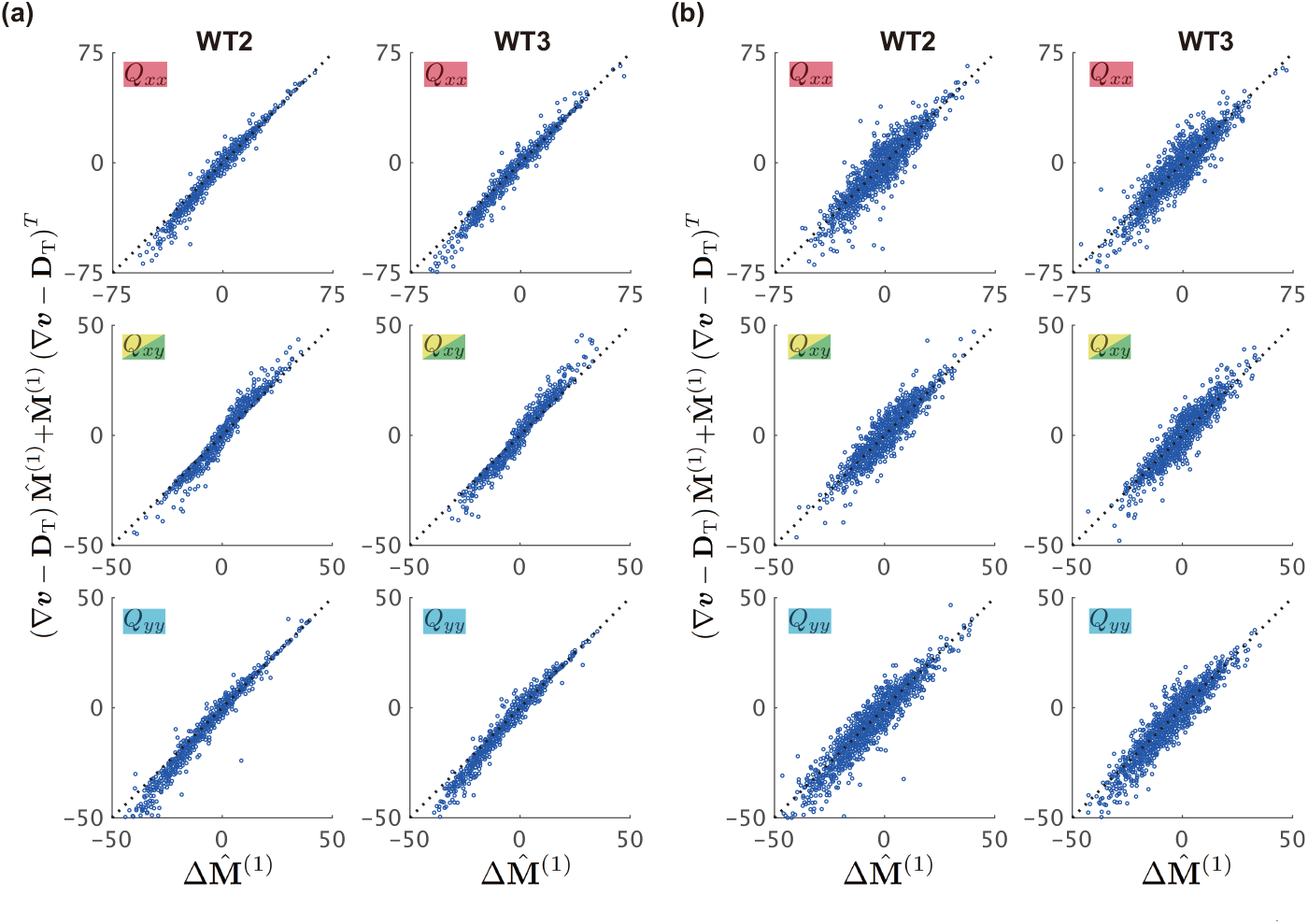
Additional data for validation of the kinematic equation Eq. 7 using 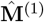. (a, b) Data from WT2 and WT3 wings (left and right columns) were utilized to calculate 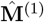 using strain rate tensors based on 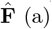 and PIV-measured ∇***v*** (b), respectively, as shown in Fig. 5b, c.

**Figure S9.**
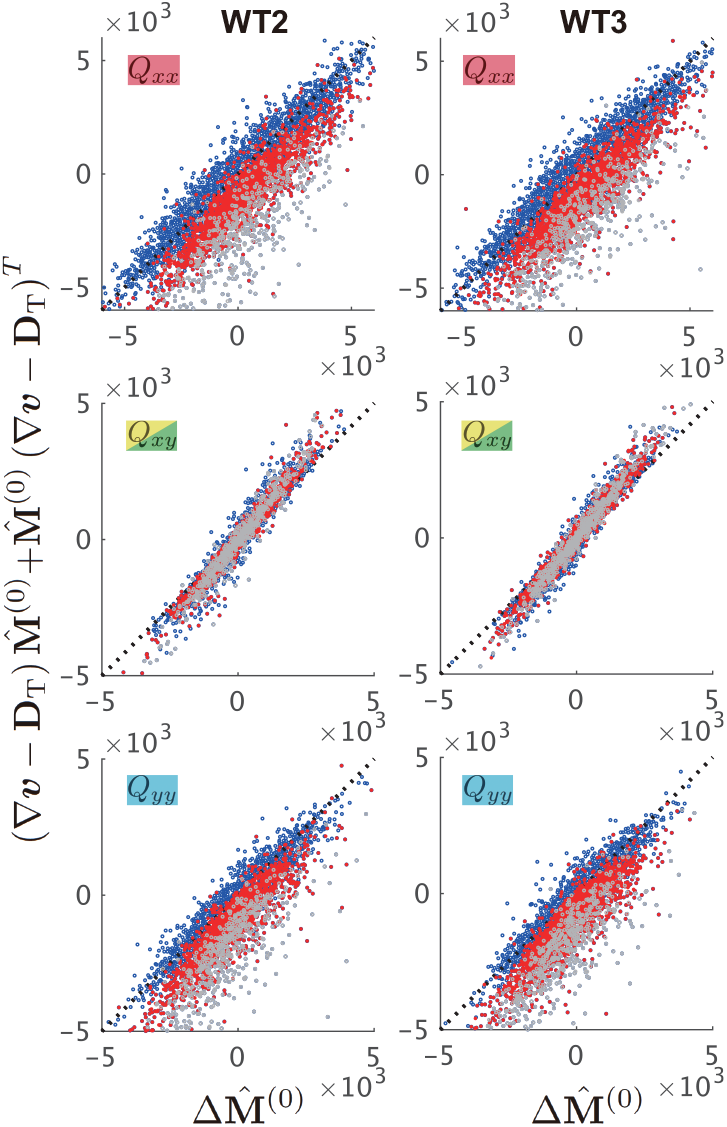
Additional data for the validation of the kinematic equation utilizing 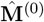. Data from WT2 and WT3 wings (left and right columns) were evaluated for the definitions in 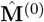 similarly to that in Fig. 5d. These point colors indicate the frequency of cell division― once (red) or multiple times (gray), or none (blue) within the ROI.

**Figure S10.**
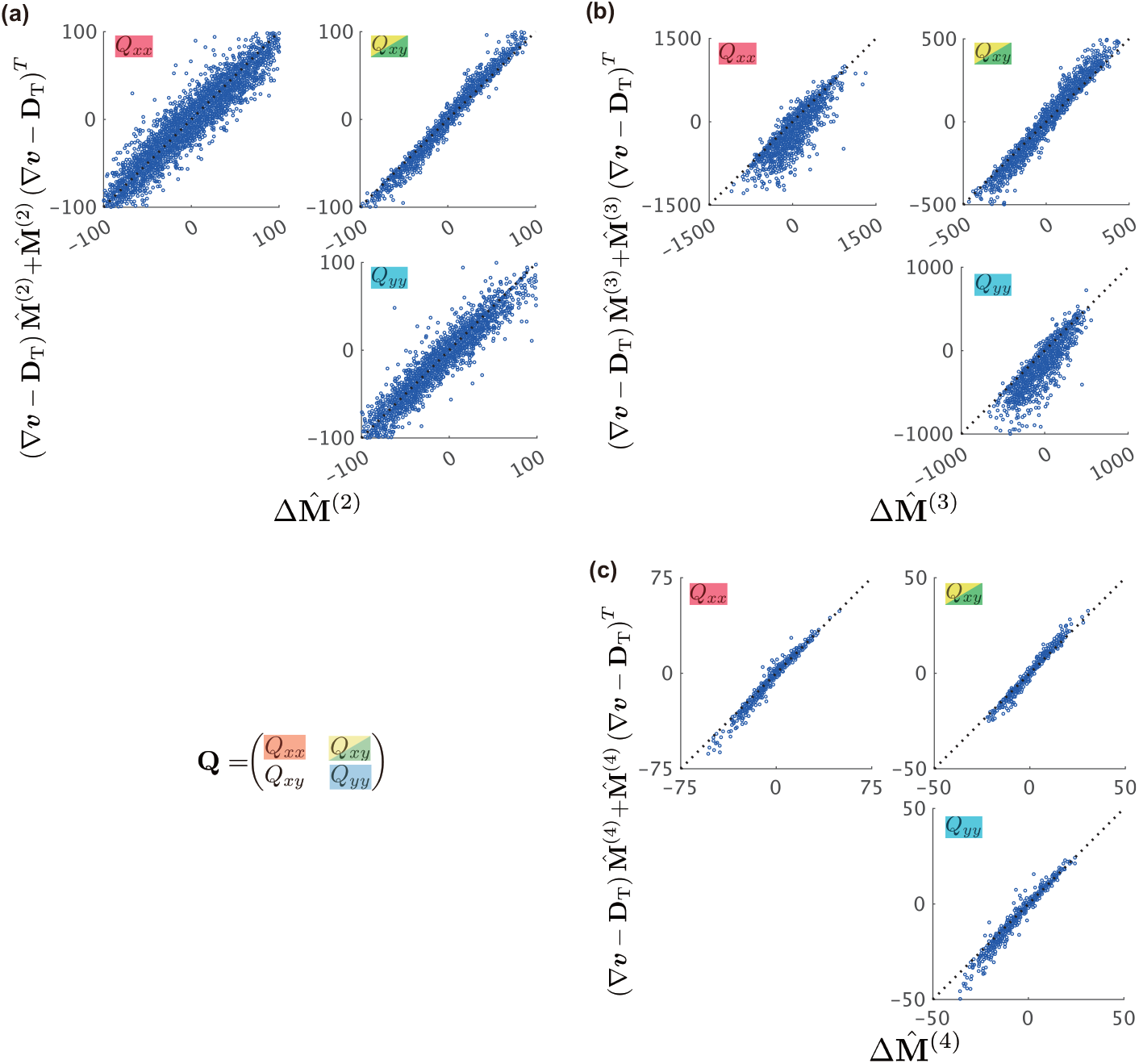
Test of the kinematic equation for different definitions of 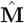. (a–c) The components of each tensor are evaluated for the following definitions of the texture tensor: (a) 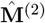, (b)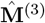, and (c)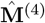. The same whole-wing data (WT1) was utilized, as in Fig. 5b, with data plotted similarly as in Fig. 5b.

**Figure S11.**
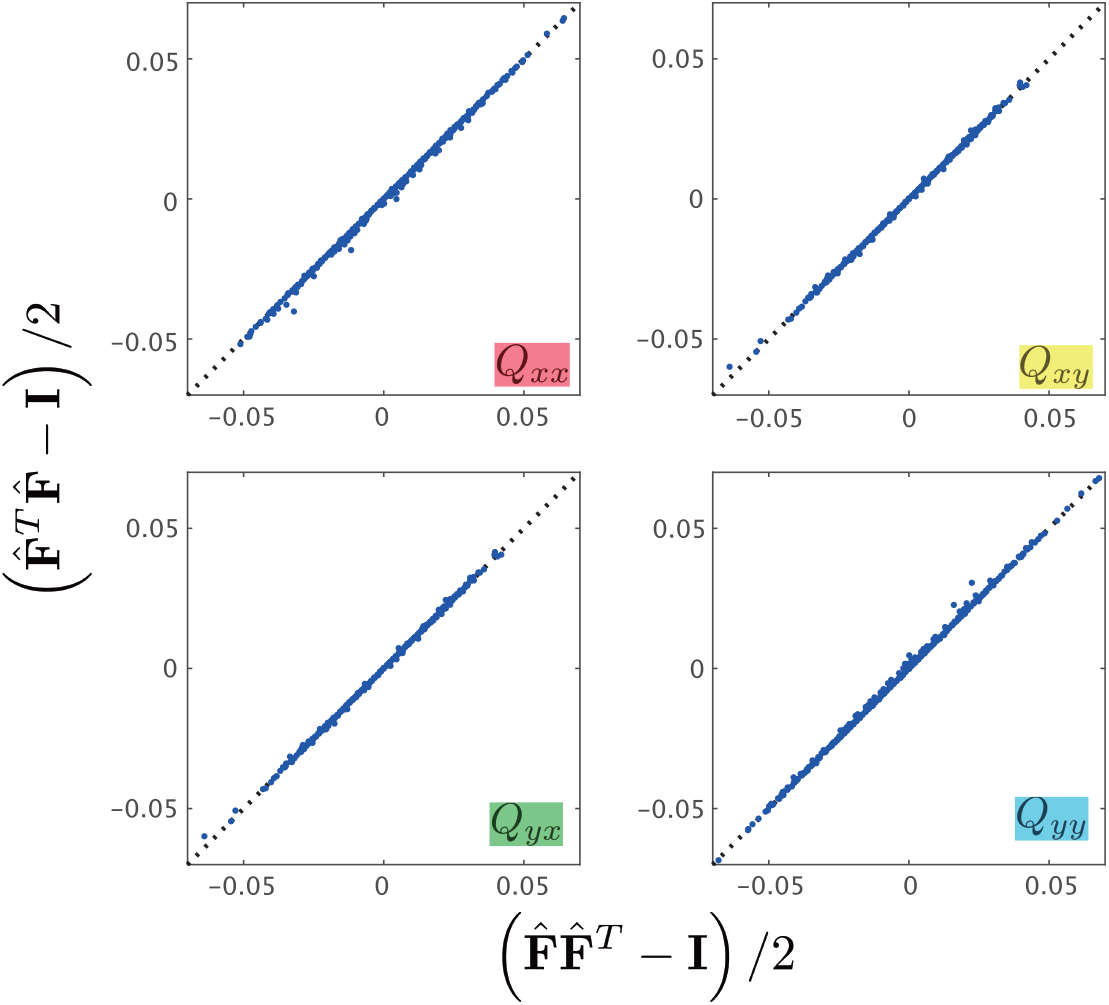
Comparison of the different formulations of the Green-Lagrange strain tensor Ê. The Green-Lagrange strain tensor utilized in this study (vertical axis) was plotted against that utilized in ref. [1] (horizontal axis).

## Footnotes

1 Φ_1_(**F**) expressed as 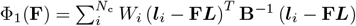, using the left Cauchy-Green tensor **B** ≡ **FF**^*T*^ . In studies on continuum mechanics, **B**^−1^ is interpreted as the metric tensor of the ***x***-space [3].

